# Assortative mating and the dynamical decoupling of genetic admixture levels from phenotypes that differ between source populations

**DOI:** 10.1101/773663

**Authors:** Jaehee Kim, Michael D. Edge, Amy Goldberg, Noah A. Rosenberg

## Abstract

Source populations for an admixed population can possess distinct patterns of genotype and phenotype at the beginning of the admixture process. Such differences are sometimes taken to serve as markers of ancestry—that is, phenotypes that are initially associated with the ancestral background in one source population are taken to reflect ancestry in that population. Examples exist, however, in which genotypes or phenotypes initially associated with ancestry in one source population have decoupled from overall admixture levels, so that they no longer serve as proxies for genetic ancestry. We develop a mechanistic model for describing the joint dynamics of admixture levels and phenotype distributions in an admixed population. The approach includes a quantitative-genetic model that relates a phenotype to underlying loci that affect its trait value. We consider three forms of mating. First, individuals might assort in a manner that is independent of the overall genetic admixture level. Second, individuals might assort by a quantitative phenotype that is initially correlated with the genetic admixture level. Third, individuals might assort by the genetic admixture level itself. Under the model, we explore the relationship between genetic admixture level and phenotype over time, studying the effect on this relationship of the genetic architecture of the phenotype. We find that the decoupling of genetic ancestry and phenotype can occur surprisingly quickly, especially if the phenotype is driven by a small number of loci. We also find that positive assortative mating attenuates the process of dissociation in relation to a scenario in which mating is random with respect to genetic admixture and with respect to phenotype. The mechanistic framework suggests that in an admixed population, a trait that initially differed between source populations might be a reliable proxy for ancestry for only a short time, especially if the trait is determined by relatively few loci. The results are potentially relevant in admixed human populations, in which phenotypes that have a perceived correlation with ancestry might have social significance as ancestry markers, despite declining correlations with ancestry over time.

**Author Summary:** Admixed populations are populations that descend from two or more populations that had been separated for a long time at the beginning of the admixture process. The source populations typically possess distinct patterns of genotype and phenotype. Hence, early in the admixture process, phenotypes of admixed individuals can provide information about the extent to which these individuals possess ancestry in a specific source population. To study correlations between admixture levels and phenotypes that differ between source populations, we construct a genetic and phenotypic model of the dynamical process of admixture. Under the model, we show that correlations between admixture levels and these phenotypes dissipate over time—especially if the genetic architecture of the phenotypes involves only a small number of loci, or if mating in the admixed population is random with respect to both the admixture levels and the phenotypes. The result has the implication that a trait that once reflected ancestry in a specific source population might lose this ancestry correlation. As a consequence, in human populations, after a sufficient length of time, salient phenotypes that can have social meaning as ancestry markers might no longer bear any relationship to genome-wide genetic ancestry.

## 1 Introduction

Admixed populations descend from two or more source groups that have long been separated and that likely possessed distinct patterns of genotype and phenotype at the beginning of the admixture process. Among individuals in an admixed population in the generations immediately after its founding, admixture levels from any specific source population are highly heterogeneous [1, 2]. For admixed individuals, measurements of specific genotypes and phenotypes that differ in frequency or distribution between source populations can often provide reasonable estimates of individual levels of genetic ancestry in the particular source populations [3, 4]—and for some phenotypes, such measurements might even be commonly regarded by researchers, societies, or admixed individuals themselves as proxies for overall genetic ancestry [5–7].

Examples exist, however, in which genotypes or phenotypes initially associated with ancestry in one source population are decoupled from overall admixture levels, so that they no longer serve as tight proxies for ancestry [5, 6, 8–13]. For example, in human genetics, consider skin pigmentation and eye color, observable traits for which the phenotypic distribution differs substantially between sub-Saharan African and European populations. In the Cape Verdean admixed population, descended from European and West African sources, measurements of skin pigmentation and eye color are correlated with sub-Saharan African genetic ancestry [11]. At the same time, the correlations between phenotype and ancestry are imperfect; many individuals with a high proportion of sub-Saharan African genetic ancestry have skin pigmentation and eye color traits in a range more typical of individuals with higher European genetic ancestry, and vice versa. Similar patterns of incomplete correlation with overall genetic ancestry hold for genotypes that underlie these phenotypes [11].

How does ancestry level become decoupled from genotype and phenotype in an admixed population? Parra et al. [8] proposed one scenario for this decoupling, using an example of assortative mating by a phenotype correlated with ancestry in Brazil. Parra et al. suggested that in Brazil, assortative mating is largely dependent on “color,” a phenotypic measure based to a large extent on skin pigmentation. According to this hypothesis, in a population descended from source groups with substantially different skin pigmentation distributions (say, sub-Saharan Africans and Europeans), similarity according to a phenotype correlated with genetic ancestry (say, *color*) increases the probability that a pair is a mating pair. Mating probabilities for pairs of individuals are more closely related to the phenotype than to overall sub-Saharan African or European genetic admixture levels *per se*. Whereas in the early generations of such a process, the phenotype would strongly reflect genetic ancestry, after a sufficient length of time with assortative mating by the phenotype, phenotypic variation would be maintained, but with similar genetic ancestry distributions for individuals with substantially different phenotype (Figure 1). Only at genes associated with the phenotype and their nearby linked genomic regions would genetic ancestry and the phenotype be associated.

**Figure 1:**
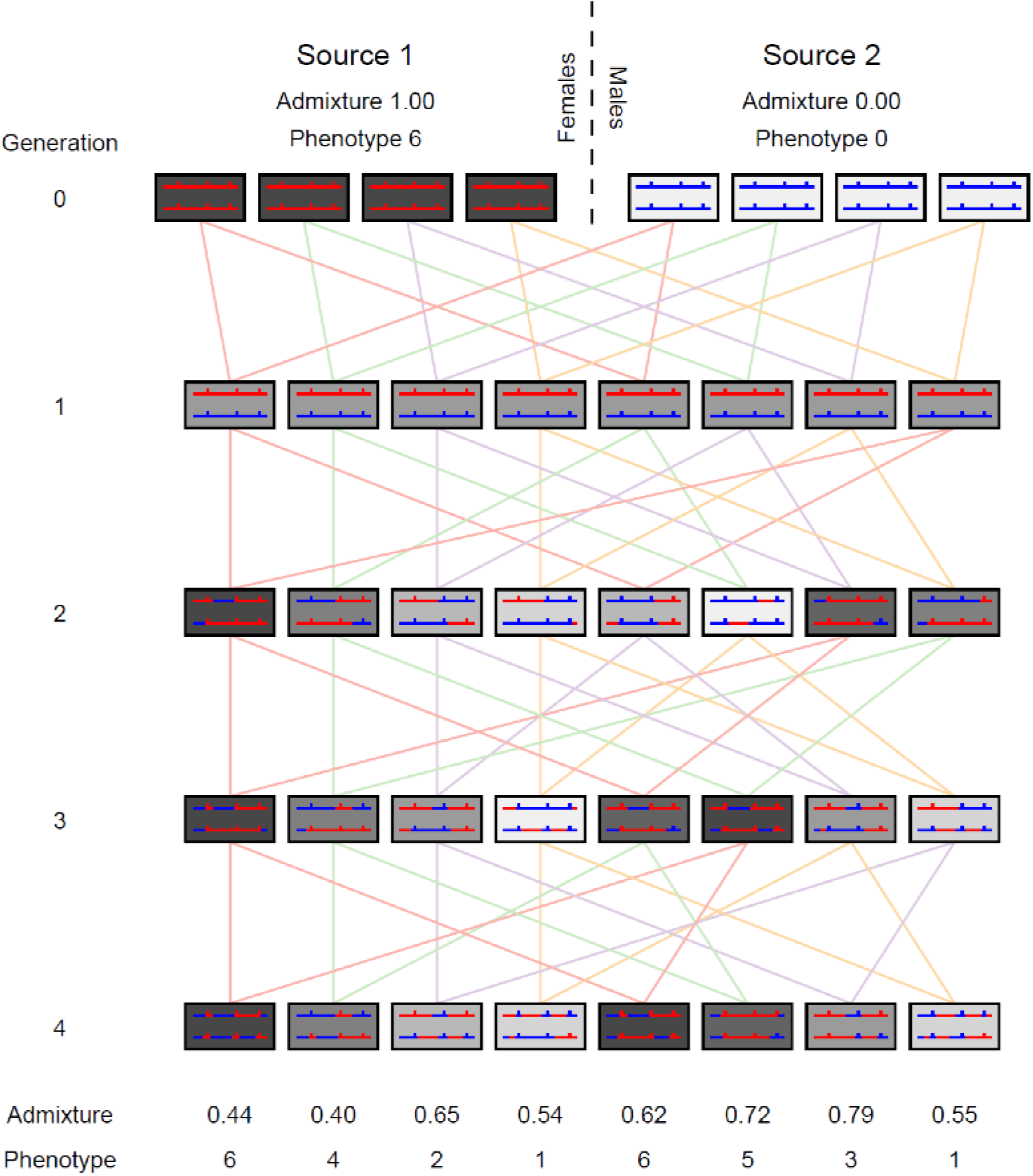
A schematic of an admixture process with positive assortative mating by a phenotype initially correlated with admixture levels. In generation 0, an admixture process begins with females from one population (source 1, left) and males from another (source 2, right). For a quantitative phenotype, source population 1 begins with a high trait value of 6 and source population 2 has a low trait value of 0. Three loci contribute additively to the genetic architecture of the phenotype; each allele derived from source population 1 contributes a value of 1 to the phenotype. The phenotype is represented by the shading of a box. Individuals are depicted as pairs of chromosomes with the ancestral sources of those chromosomes; short vertical lines along the chromosome indicate the three loci that contribute to the phenotype. After generation 1, positive assortative mating by phenotype proceeds in the admixed population. Lines connecting generations are displayed in four colors, representing four mating pairs. Initially, in generation 2, a strong correlation exists between admixture and phenotype (*r* = 0.96). By generation 4, however, owing to recombination events that stochastically dissociate the trait loci from the overall genetic admixture, the genetic admixture has been decoupled from the phenotype, so that some of the individuals with the highest trait values have among the lowest admixture coefficients for source population 1, and the correlation between phenotype and overall genetic admixture has dissipated (*r* = −0.09).

Could genetic ancestry in an admixed population become almost entirely decoupled from the phenotypes that differ between its source populations? This scenario is intriguing, as it would eliminate any connection between visible phenotypic markers of genetic ancestry and the genetic ancestry itself; the phenotype of an individual on a trait such as skin pigmentation would reveal little information about the genetic ancestry of molecular characters in an individual—other than for skin pigmentation genes and their closest genomic neighbors—nor about the total genomic ancestry of the individual.

We develop a mechanistic model describing the joint dynamics of admixture levels and phenotype distributions in an admixed population. The approach includes a quantitative-genetic model that relates a phenotype to underlying loci that affect its trait value. We consider three forms of mating. First, individuals might mate randomly, or assort independently of the overall genetic admixture level. Second, individuals might assort by a phenotype initially correlated with the genetic admixture level, but that is not identical to it. Third, following studies that have detected evidence of assortative mating by genetic admixture levels or phenotypes that are tightly connected to them [14, 15], individuals might assort by the genetic admixture level itself. Under the model, we explore the relationship between genetic admixture level and phenotype over time, studying the effect of the genetic architecture of the phenotype. We find that the decoupling of genetic ancestry and phenotype is not only possible, but it can occur surprisingly quickly, especially if the quantitative phenotype is driven by a small number of loci. Moreover, assortative mating is not required for genotype and phenotype to become decoupled. Indeed, assortative mating attenuates the process compared with a scenario in which mating is random with respect to admixture and with respect to phenotype.

## 2 Model

### 2.1 Population Model

Our mechanistic admixture model closely follows the model of Verdu, Goldberg, and Rosenberg [1, 16, 17], building on earlier related models [18–20]. We start with individuals in each of two isolated source populations, *S*_1_ and *S*_2_. At the founding of an admixed population (*g* = 0), a founding parental pool 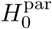 is formed, containing fraction *s*_1,0_ from population *S*_1_ and *s*_2,0_ from population *S*_2_. That is, a random individual in 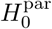 originates from population *S*_1_ with probability *s*_1,0_ and from *S*_2_ with probability *s*_2,0_. This choice requires *s*_1,0_ + *s*_2,0_ = 1 and 0 ≤ *s*_1,0_, *s*_2,0_ ≤ 1. The individuals in the founding parental pool mate according to a mating model (Section 2.3) and produce generation *g* = 1 of admixed offspring (*H*_1_).

In subsequent generations (*g* ≥ 1), in forming an admixed population *H*_*g*+1_ at generation *g* + 1, three populations contribute to its parental pool 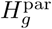: the source populations (*S*_1_ and *S*_2_) and the admixed population (*H*_*g*_) of the previous generation, with fractional contributions *s*_1,*g*_, *s*_2,*g*_, and *h*_*g*_, respectively. Here, *s*_1,*g*_, *s*_2,*g*_, and *h*_*g*_ represent probabilities for a random individual in 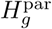 to originate from populations *S*_1_, *S*_2_, and *H*_*g*_, with constraints *s*_1,*g*_ + *s*_2,*g*_ + *h*_*g*_ = 1 and 0 ≤ *s*_1,*g*_, *s*_2,*g*_, *h*_*g*_ ≤ 1. Offspring resulting from mating among individuals in the parental pool 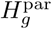 define the admixed population *H*_*g*+1_. A schematic of the admixture model appears in Figure 2.

**Figure 2:**
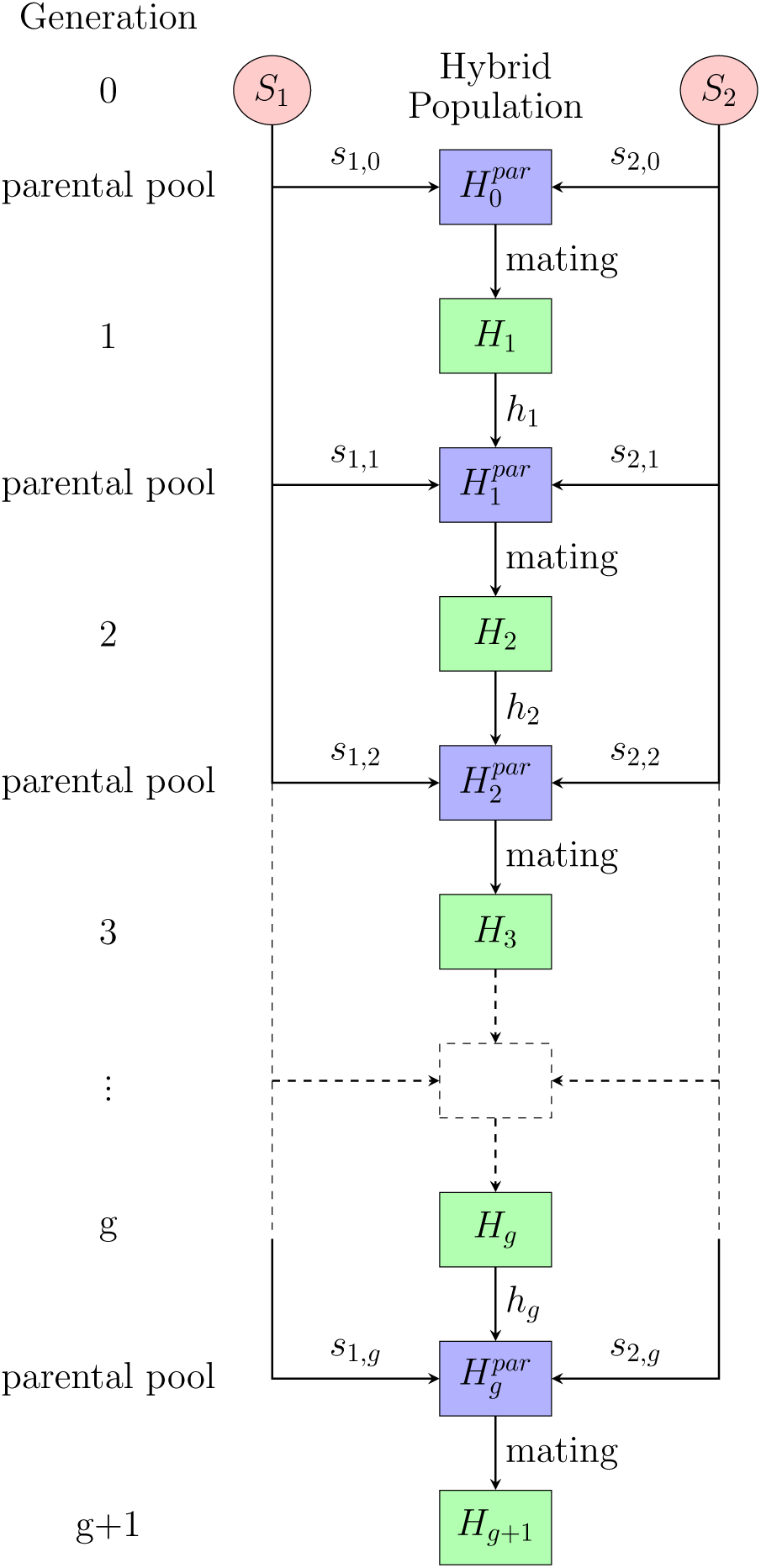
A schematic diagram of the admixture process. At the founding of the population (*g* = 0), two isolated source populations produce the first generation of a admixed population (*H*_1_). In the subsequent generations (*g* ≥ 1), populations from *S*_1_, *S*_2_, and *H*_*g*_ constitute a parental pool 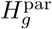 at generation *g* from which the admixed population *H*_*g*+1_ at generation *g* + 1 is produced. Fractional contributions from three populations in forming the parental pool are *s*_1,*g*_, *s*_2,*g*_, and *h*_*g*_, respectively. Individuals in the parental pool mate based on mating models described in Section 2.3.

The total admixture fraction represents the proportion of the genome of an individual originating from a specific ancestral population, *S*_1_ or *S*_2_. We denote an individual’s admixture fraction from source population *S*_1_ at generation *g* by *H*_*A,g*_, with the *A* indicating consideration of autosomal genetic loci. Given a pair of individuals with admixture fractions 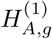 and 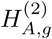, the ancestry of their offspring is deterministically set to the mean of the admixture fractions of the parents: 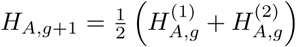. The possible values for the admixture fraction at generation *g* are 0, 1*/*2^*g*^, 2*/*2^*g*^, *…*, (2^*g*^ − 1)*/*2^*g*^, 1.

### 2.2 Quantitative Trait Model

To model a phenotype, we adopt the quantitative trait model of Edge and Rosenberg [21, 22]. We assume each individual is diploid and that *k* biallelic autosomal loci, each with the same effect size, additively determine the value of a quantitative trait. At each trait locus, we denote the allelic type more prevalent in population *S*_1_ than in population *S*_2_ as allelic type “1”, and the other allelic type as “0”. The choice is arbitrary in case the allele frequency is the same in the two populations. A diploid individual’s genotype at locus *i, i* ∈ {1, 2, *…, k*}, and allele *j, j* ∈ {1, 2}, is represented by a random indicator variable *L*_*ij*_:

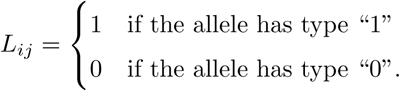

Let *M* be a random variable representing an individual’s population membership, considering individuals only from the source populations *S*_1_ and *S*_2_, and define allele frequencies for allelic type “1” at each locus given the population membership:

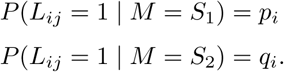

Here, *j* can be either 1 or 2. Because we define allelic type “1” to be more common in population *S*_1_ than in population *S*_2_, 0 ≤ *q*_*i*_ ≤ *p*_*i*_ ≤ 1.

An individual’s trait value is determined by a sum of contributions across loci. At each locus, we denote an allele that increases the trait value by “+” and the other allele by “−”. The total quantitative trait value *T* of an individual given a multilocus genotype is an individual’s total number of “+” alleles. Whether the “1” allelic type or “0” type is the “+” allele at locus *i* is determined by a random variable *X*_*i*_, following Edge and Rosenberg [21, 22]:

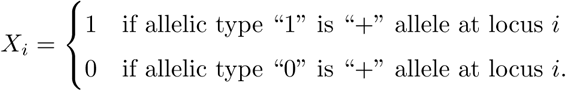

For a given set of values {*X*_1_, *X*_2_, *· · ·, X*_*k*_} for *k* quantitative trait loci, the total trait value for a diploid individual is equal to the total number of “+” alleles carried by the individuals, or:

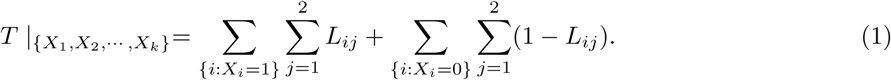

This quantity takes values in {0, 1, *…*, 2*k*}. An example of the quantitative trait model appears in Figure 3.

**Figure 3:**
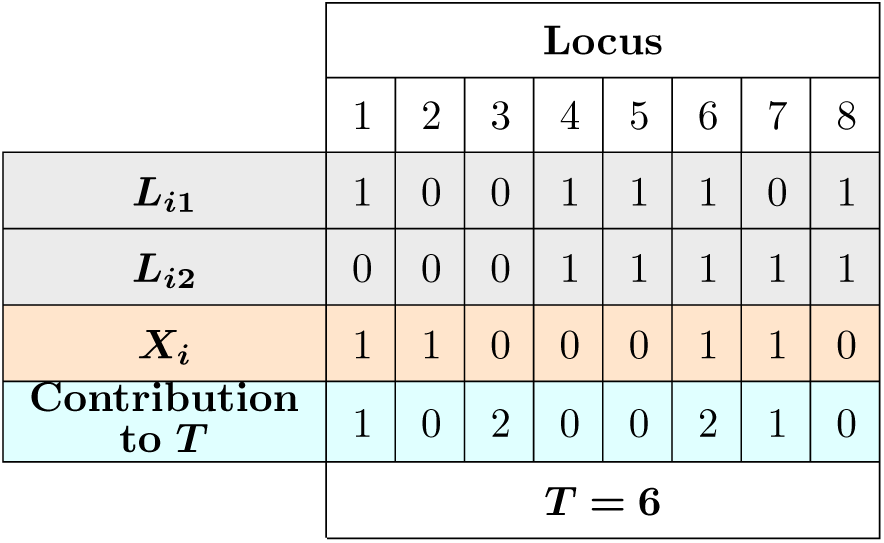
An example of our quantitative trait model. Here, a diploid individual with *k* = 8 trait loci is shown. At each locus *i*, an allele *L*_*ij*_ contributes to the overall trait value if and only if *L*_*ij*_ = *X*_*i*_ where *X*_*i*_ is a variable indicating which of two alleles, “0” or “1” increases the trait value. The total trait value of an individual equals the number of alleles satisfying *L*_*ij*_ = *X*_*i*_ across the *k* trait loci. In this example, the individual has *T* = 6.

We adopt two scenarios for the *X*_*i*_. We first consider an idealized case in which the number of “1” alleles is perfectly correlated with the trait value: *P* (*X*_*i*_ = 1) = 1 and *P* (*X*_*i*_ = 0) = 0 for all *i* = 1, 2, *…, k*, so that allelic type “1” is the “+” allele and allelic type “0” is the “−” allele for all loci. Because we define “1” to be the more frequent allelic type in source population *S*_1_, individuals from *S*_1_ are more likely to have a larger trait value than are individuals from *S*_2_. This scenario considers a case in which the phenotype is systematically different between populations 1 and 2, and is depicted in the diagram in Figure 1.

The second case is detailed in Edge and Rosenberg [21, 22], where the trait is selectively neutral during the population divergence and neither allele is preferentially correlated with the trait allele: 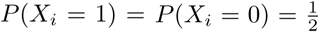 for all *i* = 1, 2, *…, k*, so that at each locus, allelic type “1” and allelic type “0” have equal probability of being the “+” allele.

### 2.3 Mating Model

We next describe three different mating models: (1) random mating, in which any two individuals have the same chance of reproducing, independent of phenotype or ancestry; (2) assortative mating by ancestry, in which the probability that two individuals reproduce depends on their ancestries; and (3) assortative mating by phenotype, in which the probability that two individuals reproduce depends on their trait values.

In each generation *g*, the parental pool 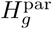 contains 2*N* individuals, *N* female and *N* male. The admixture fraction from source population *S*_1_ and trait value of a female individual *i* are denoted by 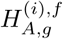 and 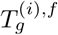, respectively. Analogous quantities for a male *j* are 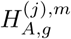 and 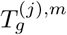. We construct an *N* × *N* mating probability matrix *M*, whose entry *m*_*ij*_ represents the probability that a female *i* and a male *j* mate:

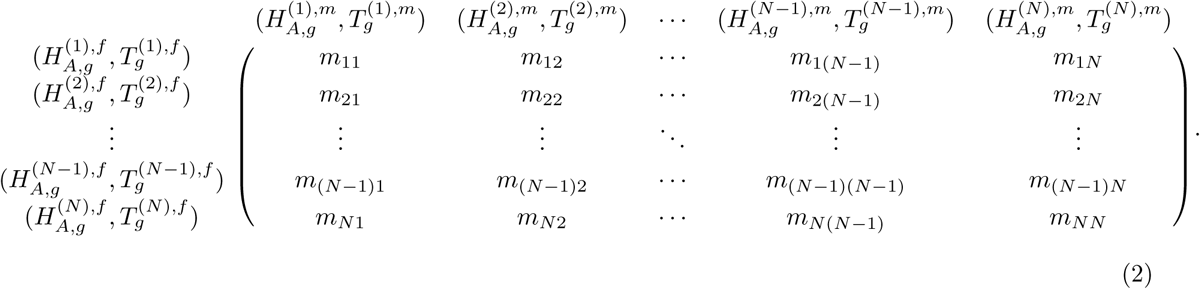

In the absence of selection, every individual in the population must have the same expected number of offspring irrespective of ancestry or phenotype. We assume that the expected number of offspring of an individual is proportional to the expected number of matings of the individual. This quantity is the sum of mating probabilities across all mates available for an individual. Therefore, the equal-offspring requirement translates into an assumption of equal row sums for females and equal column sums for males in the mating matrix in Eq. 2. Note that this assumption of equal numbers of offspring independent of ancestry and phenotype accords with a standard property of assortative mating models that assortative mating on its own does not alter allele frequencies over time [23–28].

The mating probability *m*_*ij*_ in Eq. 2 between female *i* and male *j* can be expressed as:

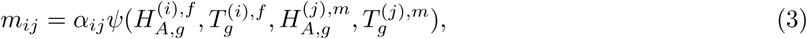

where *ψ* is a function that quantifies the dependence of the mating probability *m*_*ij*_ on the ancestry and trait values of the individuals in a pair, and *α*_*ij*_ is a normalization constant specific to the mating pair (*i, j*). The constant *α*_*ij*_ is included in order to permit the matrix entries to satisfy the constant row and column sum constraints. Without loss of generality, we choose the constant row and column sums to be 1 so that the mating probability matrix *M* is doubly stochastic. Procedures to evaluate the *α*_*ij*_ appear later in the section. The properties of *ψ* are determined by a mating model.

In random mating, the mating probability is independent of individual ancestry and trait values, so that 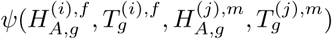 is constant across all *i* and *j*, and *m*_*ij*_ has the same value for all mating pairs. Therefore, for all pairs (*i, j*), each taken from {1, 2, *…, N*}, *m*_*ij*_ = *α*_*ij*_ = *α* for some constant *α* ∈ [0, 1].

In assortative mating by ancestry, the mating probability depends only on the ancestries of potential mates and not on the phenotypes: 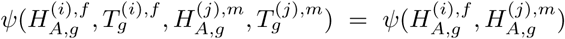 and 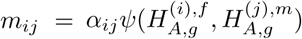. For positive assortment, the mating function has higher values if two individuals have similar ancestries and lower values as the ancestries become more different. For example, in complete assortment, *ψ* is 1 if the two input parameters have the same value and 0 if the values differ. For negative assortment, the behavior of *ψ* is reversed compared with the positive assortment case: *ψ* increases as the difference between the ancestries of the two individuals in a mating pair increases.

In assortative mating by phenotype, the mating probability depends only on the trait values of potential mates and not on the ancestries: 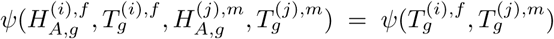 and 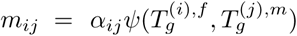. The qualitative requirements for the function *ψ* are the same as with assortative mating by ancestry, but with the trait values of the mating pair as arguments instead of the ancestries.

We adopt the following form for the mating function:

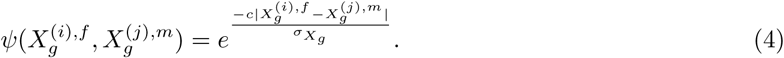

The finite constant *c* quantifies the strength of the assortative mating. For a given pair of values 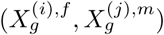, where *X*_*g*_ = *H*_*A,g*_ or *X*_*g*_ = *T*_*g*_, a larger *c* value results in a lower mating probability, which gives stronger positive assortative mating compared with a smaller *c* value. For a positive value of *c*, the function takes a value of 1 if two potential mates have the same ancestry level (or phenotype), and *ψ* decreases exponentially as the difference between the two individuals increases. A negative value of *c* indicates negative assortative mating, where two individuals with different ancestry (or phenotype) have a higher probability of mating than do two individuals with similar ancestry. We focus on positive assortative mating.

At each generation *g*, the admixture fraction *H*_*A,g*_ takes values in {0, 1*/*2^*g*^, 2*/*2^*g*^, *…*, (2^*g*^ − 1)*/*2^*g*^, 1} (Section 2.1), and the phenotype *T*_*g*_ takes values in {0, 1, *…*, 2*k*} (Section 2.2). To compare statistics from different mating schemes, we consider variables that are standardized by dividing *X*_*g*_ (*H*_*A,g*_ or *T*_*g*_) by its standard deviation 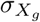 based on its distribution in 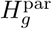 at each generation *g*. For the unstandardized variables, because *T*_*g*_ takes a higher value than *H*_*A,g*_, the effect of assortative mating by phenotype at the same assortative mating strength *c* is artificially inflated compared to the effect of assortative mating by admixture fraction.

Having specified the mating function *ψ*, we now formally state the normalization condition for the mating matrix *M*: the sum across potential mates of the mating probabilities of a random individual in the parental pool must be 1. Recalling that each entry *m*_*ij*_ in *M* represents the probability that individuals *i* and *j* mate, the condition requires the row and column sums in the mating matrix to be 1 for each row and column. We start with an unnormalized mating matrix 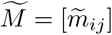 whose entries are:

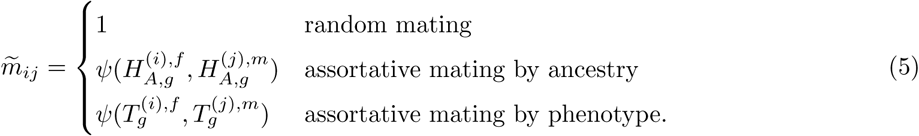

We must obtain *N*^2^ normalizing constants *α*_*ij*_ such that the mating matrix 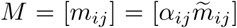 satisfies the double stochasticity requirement. This requirement gives 2*N* constraints, one for each row and one for each column. Because each entry in the mating matrix represents the probability for two individuals to mate, we also require *m*_*ij*_ to be in [0, 1] for all (*i, j*) ∈ {1, 2, *…, N*}^2^.

Infinitely many matrices satisfy the constraints, as the set of 2*N* equations with *N*^2^ variables is underdetermined. We choose the matrix *M* by identifying the matrix that satisfies the set of constraints and that is closest to our model matrix 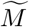 according to the principle of minimum discrimination information (pp. 36-43 in [29]). Here, the “closeness” of a pair of matrices is measured by the Kullback-Leibler divergence *D*_*KL*_ (pp. 1-11 in [29]), which is nonnegative and is equal to zero if and only if the two matrices are identical.

The problem of identifying *M* can be formally written as a convex optimization problem. The objective function that we seek to minimize is

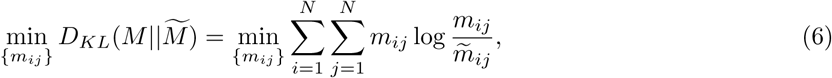

and we have constraints

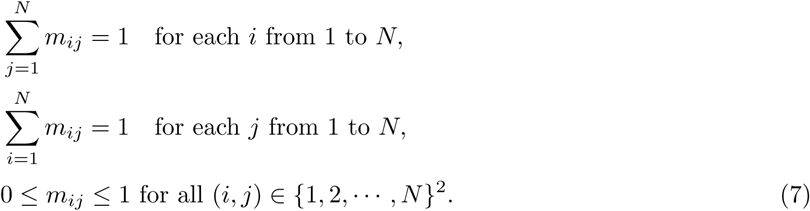

We use the interior-point method [30, 31], which iteratively traverses within the feasible region to obtain the optimal solution numerically, as implemented in `mosek` function of R package `Rmosek` [32]. For fixed 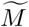, the Hessian of the KL divergence has

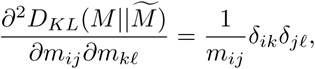

where *δ* is the Kronecker delta. Because ∇^2^*D*_*KL*_ > 0 for all *m*_*ij*_ ∈ (0, 1), the KL divergence function is strictly convex (Section 3.1.4 in [33]) in each of the *N* ^2^ variables in *M* for fixed 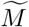, and thus, the optimal solution found by numerical minimization is the unique global minimum (Section 4.2.1 in [33]).

### 2.4 Expectation and Variance of the Admixture Fraction

To interpret our simulations of admixture dynamics, we will need a series of results concerning the mean and variance of the admixture fraction in the admixed population. In particular, we derive a relationship between the variance of the admixture fraction and the correlation in admixture levels for members of mating pairs.

Let *H*_*A,g*_ be a random variable representing the admixture fraction of an individual chosen at random in the admixed population *H*_*g*_ at generation *g* ≥ 1. We denote by 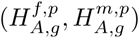 the admixture fractions of the members of a mating pair chosen at random from a parental pool 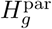 in generation *g* ≥ 0. Here, the superscript *p* denotes that the individual is from the parental pool. The parental pool 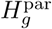, from which the admixed population *H*_*g*+1_ at generation *g* + 1 is formed, consists of populations *S*_1_, *S*_2_, and *H*_*g*_, with fractional contributions *s*_1,*g*_, *s*_2,*g*_, and *h*_*g*_ respectively (Section 2.1 and Figure 2). Because we assume each population (*S*_1_, *S*_2_, and *H*_*g*_) has equally many males and females, the numbers of males and females each remain constant at *N*, every individual has the same expected number of offspring, and no sex bias by population of origin exists in parental pairings (Section 2.3), 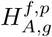 and 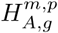 are identically distributed.

Let the random variable *Y* indicate the population membership of a random individual in 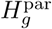. Then

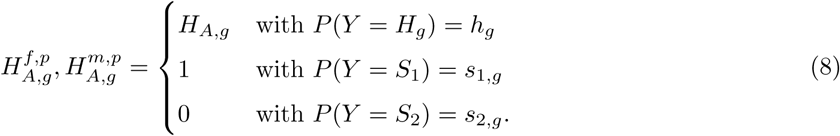

For the expectation of admixture in the parental pool, we have

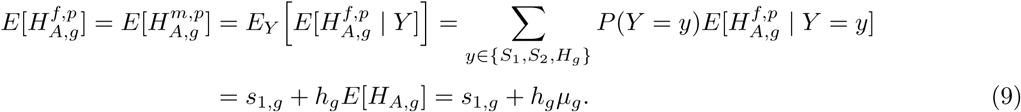

As a consequence of Eq. 9, we also have

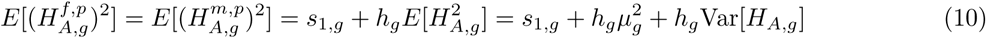

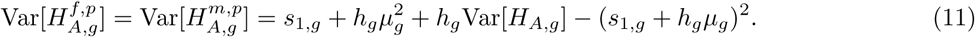

Here, *µ*_*g*_ = *E*[*H*_*A,g*_] indicates the expectation of the admixture fraction of a random individual in the admixed population *H*_*g*_ at generation *g* ≥ 1.

The ancestry of an offspring individual is deterministically set to the mean of the admixture fractions of the parents. This choice gives:

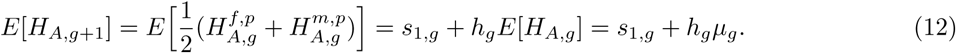

We obtain the recursion for the variance of the admixture fraction over a single generation as follows:

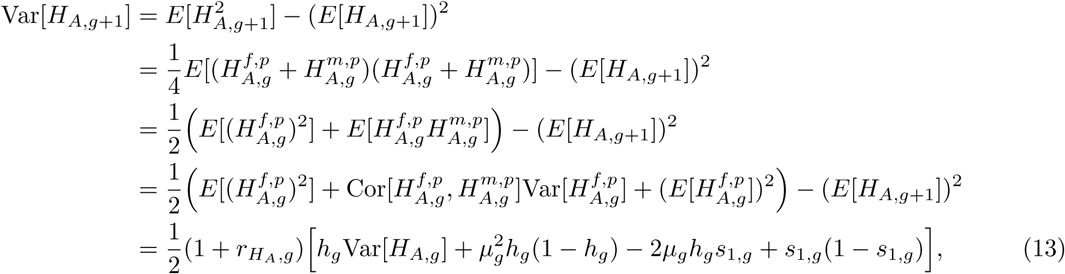

where 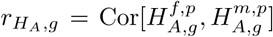 denotes the correlation of the admixture fractions in a mating pair. The last step is obtained from Eqs. 9–12. As we will see, the time-varying 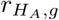 value in general depends on the parameters of the population model, the quantitative trait model, and the mating model.

For a special case of a single admixture event in which source populations *S*_1_ and *S*_2_ do not contribute to the admixed population after its founding (*s*_1,*g*_ = *s*_2,*g*_ = 0 and *h*_*g*_ = 1 for all *g* ≥ 1), the expectation of the admixture fraction stays constant in time (Eq. 12), and the variance reduces to a simple formula:

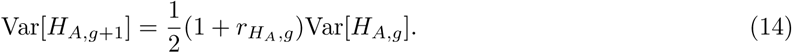

Under random mating in an infinite population with no ongoing contributions from the source populations, with 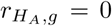 for all *g* ≥ 0, Eq. 14 reduces to the formula Var[*H*_*A,g*_] = *s*_1,0_(1 − *s*_1,0_)*/*2*g* of Verdu and Rosenberg [1]. Eq. 14 has also been derived by Zaitlen et al. [27] but under assumptions of constant mating correlation 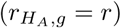 across all generations, no migration, and infinite population size.

## 3 Simulation

### 3.1 Simulation Procedure

Having specified the populations of interest, the properties of trait values in the populations, and the mating probabilities for pairs of individuals, we now describe how we simulate populations under the model. At the first time step (*g* = 0), *s*_1,0_*N* and *s*_2,0_*N* males are randomly generated from the source populations *S*_1_ and *S*_2_, respectively, with *s*_1,0_ + *s*_2,0_ = 1. The corresponding numbers of females *s*_1,0_*N* and *s*_2,0_*N* are randomly drawn from source populations, *S*_1_ and *S*_2_, respectively, constituting the founding parental pool 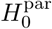 of 2*N* individuals, with *N* males and *N* females. All individuals in source population *S*_1_ have an admixture fraction value of 1, and all individuals in source population *S*_2_ have an admixture fraction value of 0, by definition. For each individual in populations *S*_1_ and *S*_2_, genotypes at each of *k* quantitative trait loci are then randomly generated on the basis of pre-specified allele frequencies *p*_*i*_ and *q*_*i*_.

We consider two different distributions for the *p*_*i*_ and *q*_*i*_. First, we assume that the two source populations display fixed differences at all trait loci, so *p*_*i*_ = 1 and *q*_*i*_ = 0 for all *k* loci. In this case, every individual in population *S*_1_ has the “1” allele at all trait loci, and every individual in population *S*_2_ has the “0” allele at all trait loci. In subsequent generations, allele “1” can be traced back to population *S*_1_, and allele “0” to *S*_2_ (Figure 1). This choice for the *p*_*i*_ and *q*_*i*_ models a case in which trait-influencing alleles are initially entirely predictive of ancestry and vice versa.

Second, departing from the idealized model, we simulate sets of *k* allele frequency pairs (*p*_*i*_, *q*_*i*_), *i* ∈ {1, *…, k*} following Edge and Rosenberg [22]. Allele frequencies *π*_*i*_ for derived alleles in the “ancestral” population of *S*_1_ and *S*_2_ are drawn based on the neutral site frequency spectrum: *P* [*π*_*i*_ = *j/*(2*N*_*a*_)] ∝ 1*/j*, where *N*_*a*_ indicates the size of the ancestral population (Eq. B6.6.1 in [34]). We use 2*N*_*a*_ = 20, 000. We assume each locus *i* in *S*_1_ and *S*_2_ undergoes independent genetic drift following a split. We add random numbers *ϵ*_*i*,1_ and *ϵ*_*i*,2_ drawn from a Normal(0, *γπ*_*i*_(1 − *π*_*i*_)) distribution to *π*_*i*_ to simulate derived allele frequencies at locus *i* in populations *S*_1_ and *S*_2_, respectively. The parameter *γ* represents the amount of variance introduced by drift into the allele frequencies of the divergent populations. Following Edge and Rosenberg [22], we choose *γ* = 0.3 so that the overall degree of genetic differentiation between *S*_1_ and *S*_2_ at a group of simulated loci approximates worldwide human *F*_*ST*_ estimates. If *ϵ*_*i*,1_ ≥ *ϵ*_*i*,2_, then we assign *p*_*i*_ = *π*_*i*_ + *ϵ*_*i*,1_ and *q*_*i*_ = *π*_*i*_ + *ϵ*_*i*,2_. If *ϵ*_*i*,1_ *< ϵ*_*i*,2_, then we assign *p*_*i*_ = 1 − (*π*_*i*_ + *ϵ*_*i*,1_) and *q*_*i*_ = 1 − (*π*_*i*_ + *ϵ*_*i*,2_). Note that if this procedure produces *p*_*i*_ > 1 or *q*_*i*_ *<* 0, then we assign *p*_*i*_ = 1 and *q*_*i*_ = 0 so that 0 ≤ *q*_*i*_ ≤ *p*_*i*_ ≤ 1.

Using the mating function in Eq. 4, we compute an unnormalized mating matrix for every pair containing a male and a female from the parental pool:

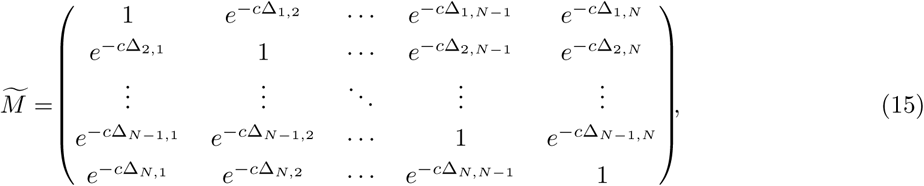

where 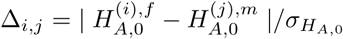 for assortative mating by ancestry and 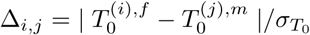 for assortative mating by trait. For random mating, *c* = 0 and all entries equal 1. The matrix 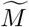 is normalized using the procedure of Section 2.3, producing the mating probability matrix *M*.

Considering all *N*^2^ potential mating pairs, we randomly draw *N* mating pairs with replacement from the parental pool, weighting mate choices by the mating probabilities in *M*. Once mates are chosen, each mating pair produces two children, one male and one female, in order to keep the population size of the offspring generation constant at *N* males and *N* females. An admixture fraction for an offspring individual is then assigned as the mean of its parental admixture fractions. Assuming no linkage disequilibrium and no mutation, the genotype of the offspring at the quantitative trait loci is then determined by independently selecting at each locus one random allele from one parent and one from the other. The resulting 2*N* offspring form the admixed population *H*_1_ at generation 1.

In subsequent generations *g* ≥ 1, we randomly select *s*_1,*g*_*N, s*_2,*g*_*N*, and *h*_*g*_*N* males and *s*_1,*g*_*N, s*_2,*g*_*N*, and *h*_*g*_*N* females from *S*_1_, *S*_2_, and *H*_*g*_, respectively, forming a *g*th generation parental pool 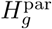 of 2*N* individuals, consisting of *N* males and *N* females. The procedure to generate the offspring population *H*_*g*+1_ from 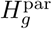 is the same as the procedure for generating *H*_1_ from 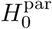.

Throughout the simulation, we keep the population size parameter *N* constant at 1,000 for computational efficiency in the matrix normalization step. Here, the admixed population size (*N*) is not necessarily identical to the source population sizes (*N*_*a*_). For each set of parameters, (*k, p*_1_, *p*_2_, *…, p*_*k*_, *q*_1_, *q*_2_, *…, q*_*k*_, *X*_1_, *X*_2_, *…, X*_*k*_, *c, s*_1,0_, *s*_1,*g*_, *s*_2,*g*_), we propagated the population to *G* = 40 generations. We generated 100 independent trajectories for each parameter set. For each trajectory, we computed statistics of interest, averaging them over all 100 trajectories in the simulation given the fixed set of parameters.

### 3.2 Base Case

We start with an idealized base case. First, we specify the parameters involving the population model (Section 2.1). We assume an equal influx from each source population at founding *g* = 0: *s*_1,0_ = 0.5, *s*_2,0_ = 1 − *s*_1,0_ = 0.5. We also assume no additional contributions from the source populations in the subsequent generations, *s*_1,*g*_ = *s*_2,*g*_ = 0, and *h*_*g*_ = 1 − *s*_1,*g*_ − *s*_2,*g*_ = 1 for all *g* ≥ 1.

Next, we choose parameter values for the quantitative trait model (Section 2.2). We consider *k* = 10 trait loci. Across the *k* loci, all “1” alleles come from source population *S*_1_ and all “0” alleles come from *S*_2_: *p*_*i*_ = 1 and *q*_*i*_ = 0 for all *i* = 1, 2, *…, k*. For each locus *i* contributing to the quantitative trait, we define “1” to be the “+” allele and “0” to be the “−” allele: *X*_*i*_ = 1 for all *i* = 1, 2, *…, k*.

Finally, for the mating model (Section 2.3), we set the assortative mating strength *c* in Eq. 4 to 0.5.

### 3.3 Statistics Measured

In each simulated admixed population, in each generation *g*, we computed the following statistics: correlation between admixture fraction and trait (Cor[*H*_*A*_, *T*]), variance of the admixture fraction in the population (Var[*H*_*A*_]), and variance of the trait value in the population (Var[*T*]). In the following section, we discuss how these statistics of interest change as we modify the simulation parameters.

## 4 Results

### 4.1 Base Case

#### 4.1.1 Correlation between genetic ancestry and phenotype (Cor[*H*_*A*_, *T*])

In the base case, each individual from *S*_1_ has admixture fraction *H*_*A*_ = 1 and trait value *T* = *k*, and each individual from *S*_2_ has admixture fraction *H*_*A*_ = 0 and trait value *T* = 0. Therefore, in the founding parental pool, 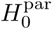, admixture fraction and trait value are perfectly correlated: Cor[*H*_*A*_, *T*] = 1. In subsequent generations, however, the correlation between the admixture fraction and the trait values starts to decouple, as illustrated in Figure 1. With all parameters involving the population model and the quantitative trait model fixed, the rate of decay in Cor[*H*_*A*_, *T*] depends on the mating model.

A comparison of Cor[*H*_*A*_, *T*] under the three mating models using base-case parameters appears in Figure 4E. Irrespective of the mating model, the founding parental pool has a perfect correlation between ancestry and phenotype. Even if the population starts with perfect correlation between admixture fractions and trait values, however, then random mating rapidly decouples them (red curve). It takes 6 generations of random mating for the correlation to decrease below 0.5 (Cor[*H*_*A*_, *T*] = 0.490). After *g* = 20 generations, the correlation becomes 0.137, and it is near zero at *g* = 40 (−0.003).

**Figure 4:**
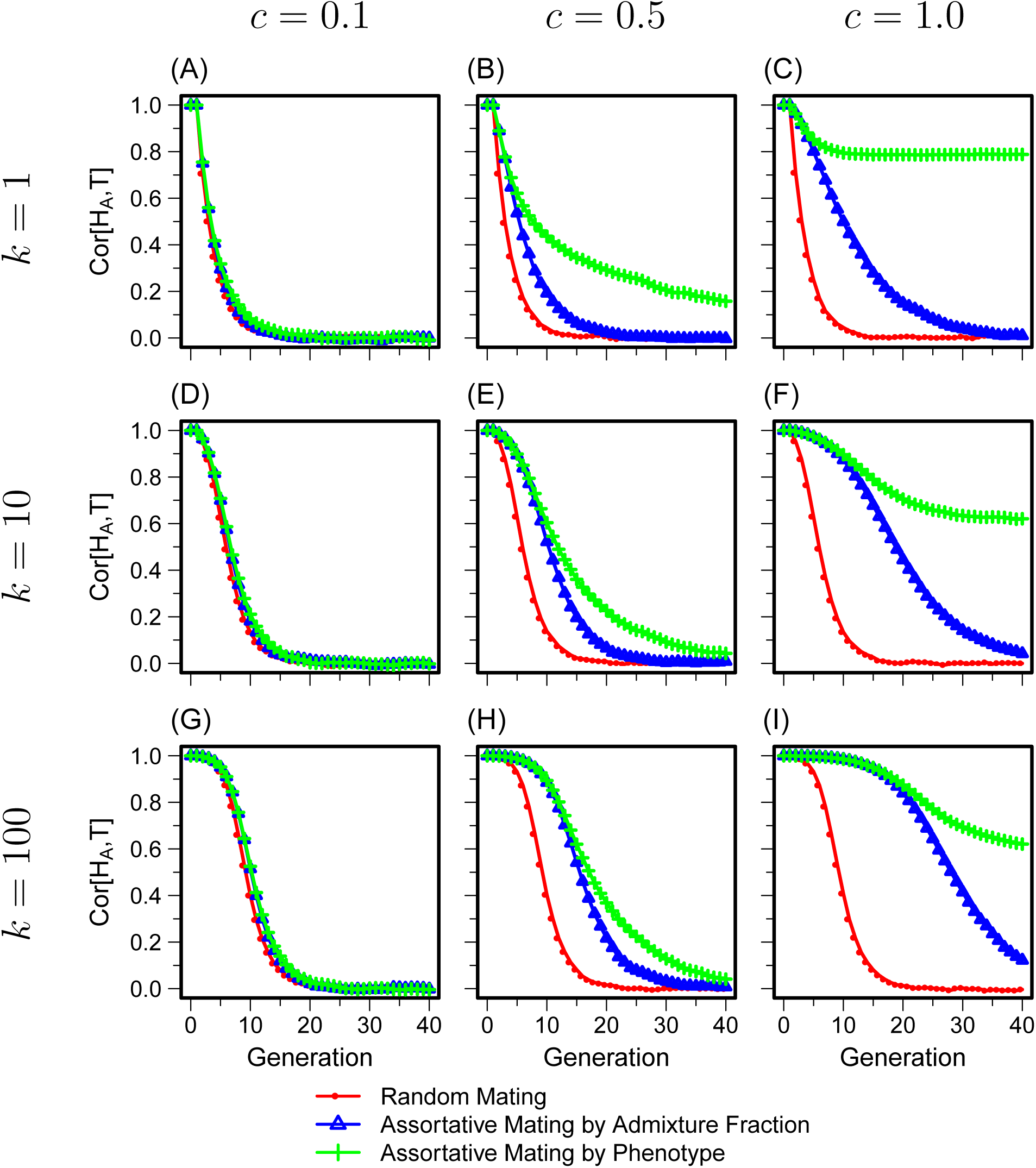
Correlation between admixture fraction and quantitative trait value (Cor[*H*_*A*_, *T*]) as a function of time. All parameter values in panel (E) follow the base case described in Section 3.2; the number of quantitative trait loci *k* and the assortative mating strength *c* vary across panels. In each panel, for a given (*k, c*) pair, for each mating scheme, the mean of 100 simulated trajectories is plotted. The red, blue, and green curves represent results from random mating, assortative mating by admixture fraction, and assortative mating by phenotype, respectively. (A) *k* = 1, *c* = 0.1. (B) *k* = 1, *c* = 0.5. (C) *k* = 1, *c* = 1.0. (D) *k* = 10, *c* = 0.1. (E) *k* = 10, *c* = 0.5. (F) *k* = 10, *c* = 1.0. (G) *k* = 100, *c* = 0.1. (H) *k* = 100, *c* = 0.5. (I) *k* = 100, *c* = 1.0.

Compared to random mating, positive assortative mating slows the decoupling of admixture fractions and trait values. Assortative mating by phenotype (green curve in Figure 4E) maintains the correlation longer than assortative mating by admixture fraction (blue curve in Figure 4E). It takes 11 generations under assortative mating by phenotype for the correlation to drop below 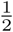 (Cor[*H*_*A*_, *T*] = 4.490), and 10 generations under assortative mating by admixture (Cor[*H*_*A*_, *T*] = 0.443). Across the 40 generations we simulated, Cor[*H*_*A*_, *T*] is consistently higher under assortative mating by phenotype than under assortative mating by admixture fraction. The correlation decreases to 0.227 at *g* = 20 and 0.043 at *g* = 40 under assortative mating by phenotype. The corresponding values under assortative mating by admixture are 0.065 at *g* = 20 and 0.009 at *g* = 40, both considerably lower than under assortative mating by phenotype.

The speed of the decoupling between admixture fraction and trait value increases with two factors: Mendelian noise in generating admixed individuals from the admixed population and decreasing contributions from the source populations. Compared to random mating, both assortative mating models have higher probabilities for matings within source populations, and thus, the proportion of individuals produced in the admixed population at *g* = 1 that are genetically admixed is smaller (blue and green lines in the marginal plots for *H*_*A*_ in Figure S1A). Over time, as displayed in Figure S1, random mating pulls individuals away from the source populations, pushing the *H*_*A*_ and *T* distributions towards the mean values rapidly. On the other hand, both assortative mating models maintain individuals with *H*_*A*_ and *T* values near the source population values for longer, and thus, they retain higher Cor[*H*_*A*_, *T*] than random mating.

The difference in Cor[*H*_*A*_, *T*] between the two assortative mating models arises from the difference between Var[*H*_*A*_] and Var[*T*]. Cov[*H*_*A*_, *T*] is similar under the two models. Given the similar covariance,

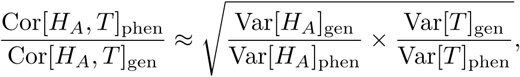

where the subscripts “gen” and “phen” indicate the property on which mating pairs assort. As we show in the next section, both assortative mating models increase Var[*H*_*A*_] and Var[*T*] compared to random mating, and the increase in variance is the largest for the property on which mating assorts: Var[*H*_*A*_]_gen_ > Var[*H*_*A*_]_phen_ and Var[*T*]_phen_ > Var[*T*]_gen_. However, we will see that the increase in Var[*H*_*A*_] due to assortative mating by admixture fraction exceeds the increase in Var[*T*] due to assortative mating by trait:

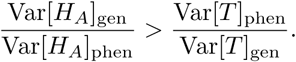

This result leads to higher Cor[*H*_*A*_, *T*] under assortative mating by trait compared to that under assortative mating by admixture fraction.

#### 4.1.2 Variance of Ancestry and Phenotype (Var[*H*_*A*_] and Var[*T*])

Each individual in *S*_1_ has admixture fraction 1, and each individual in *S*_2_ has admixture fraction 0. In the founding parental pool, Var[*H*_*A*_] = 0.250 for all three mating models. As discussed in Section 2.4, the variance of the admixture fraction can be understood in relation to the correlation coefficient 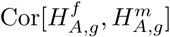 of the admixture fractions of members of mating pairs. Figure S2 shows this correlation coefficient for the simulations of Figure 4, and Figure S3 shows the analogous correlation 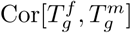 of trait values.

Figure 5E then shows the variance of the admixture fraction over time under the three mating models, for the same simulations from Figure 4E with the base case parameters. The Var[*H*_*A*_] curves in Figure 5E under the three mating models follow Eq. 13, using the time-varying 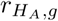 in Figure S2.

**Figure 5:**
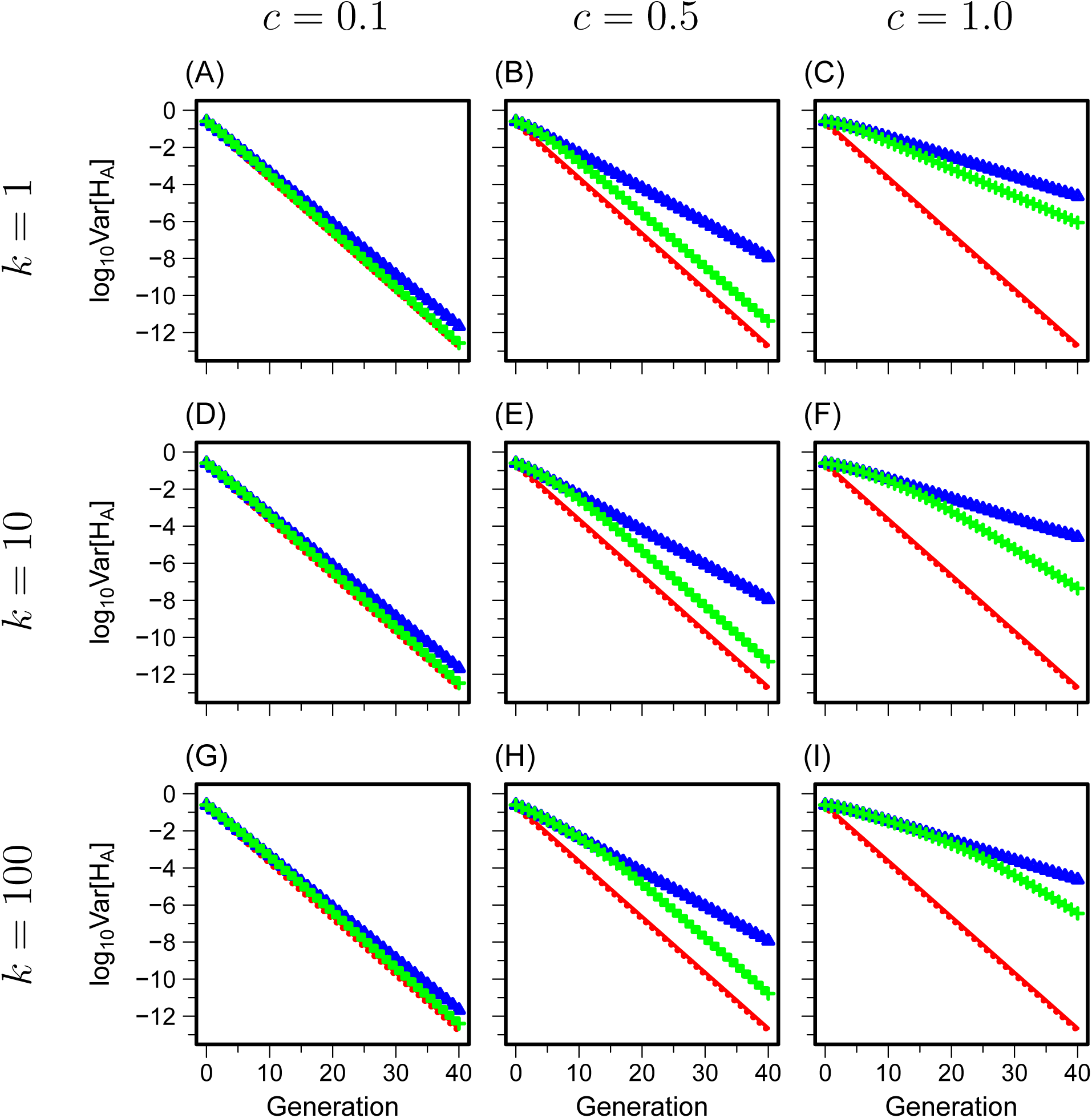
Variance of admixture fraction (Var[*H*_*A*_]) as a function of time. The simulations shown are the same ones from Figure 4. (A) *k* = 1, *c* = 0.1. (B) *k* = 1, *c* = 0.5. (C) *k* = 1, *c* = 1.0. (D) *k* = 10, *c* = 0.1. (E) *k* = 10, *c* = 0.5. (F) *k* = 10, *c* = 1.0. (G) *k* = 100, *c* = 0.1. (H) *k* = 100, *c* = 0.5. (I) *k* = 100, *c* = 1.0. Colors and symbols follow Figure 4. The y-axis is plotted on a logarithmic scale.

Among the three mating models, Var[*H*_*A*_] decreases fastest for random mating. After one generation, Var[*H*_*A*_] falls in half (0.125), and it continues to decrease monotonically by half. After 40 generations, the value decreases to 2.118 × 10^−13^. The distribution of the admixture fraction concentrates around 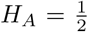 at each generation. Because the offspring admixture fraction is the mean of those of its parents, without additional influx from the source populations after the founding event, random mating rapidly drives the admixture fraction away from extreme values (0 or 1) toward the mean value of the parental pool 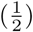.

Under assortative mating by admixture, pairs with similar admixture fraction have higher mating probabilities than under random mating. The fraction of offspring that are admixed is smaller than under random mating, and the admixture fraction distribution remains close to the extreme values (0 or 1) for longer (Figure S1). Hence, Var[*H*_*A*_] is larger under assortative mating by admixture fraction (Figure 5E). Without influx from the source populations, Var[*H*_*A*_] eventually decreases to zero, but the decrease is slower than for random mating. Var[*H*_*A*_] = 0.184 after one generation of assortative mating by admixture fraction, and Var[*H*_*A*_] = 1.118 × 10^−8^ after 40 generations. This result can also be seen in Eq. 13. From generation *g* to *g* + 1, Var[*H*_*A*_]_*g*_ decreases by a factor of 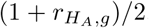. With positive assortative mating by admixture 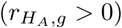, Var[*H*_*A*_] in the next generation is increased compared to the case of random mating 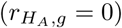.

Under assortative mating by phenotype, Var[*H*_*A*_] = 0.183 after one generation of assortative mating by phenotype, and Var[*H*_*A*_] = 4.910 × 10^−12^ after 40 generations. For the first few generations (*g <* 5), because Cor[*H*_*A*_, *T*] is high, the correlation between the admixture fraction of mating pairs, and thus Var[*H*_*A*_], is similar under the two assortative mating models, as shown in the comparison of the green and blue curves in Figures S2 and 5. However, because the admixture fraction and phenotype decouple by introduction of the Mendelian noise, mating assortatively by phenotype results in lower 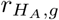 than mating assortatively by admixture fraction. In accord with Eq. 13, assortative mating by phenotype produces faster decay in Var[*H*_*A*_] with its lower 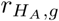 at each generation than assortative mating by admixture fraction.

For the variance of the phenotype, using Eq. 1, all individuals in *S*_1_ and *S*_2_ have trait values of 20 and 0, respectively. Therefore, in the founding parental pool, 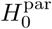, noting that *S*_1_ and *S*_2_ each have 1,000 individuals, this variance has the same constant value of 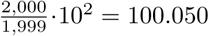 irrespective of the mating model. Figure 6E displays the variance of the phenotype, which decreases most rapidly under random mating. After one generation of random mating, Var[*T*] decreases by half (50.025), and it approaches its steady-state value of ≈ 4.957 after 13 generations. Opposite to what was seen for Var[*H*_*A*_], however, assortative mating by trait retains Var[*T*] higher for longer than assortative mating by admixture fraction. Similar to the case with Var[*H*_*A*_] under assortative mating by admixture fraction, assortative mating by phenotype keeps the trait values close to extreme values for longer than the other two mating models.

**Figure 6:**
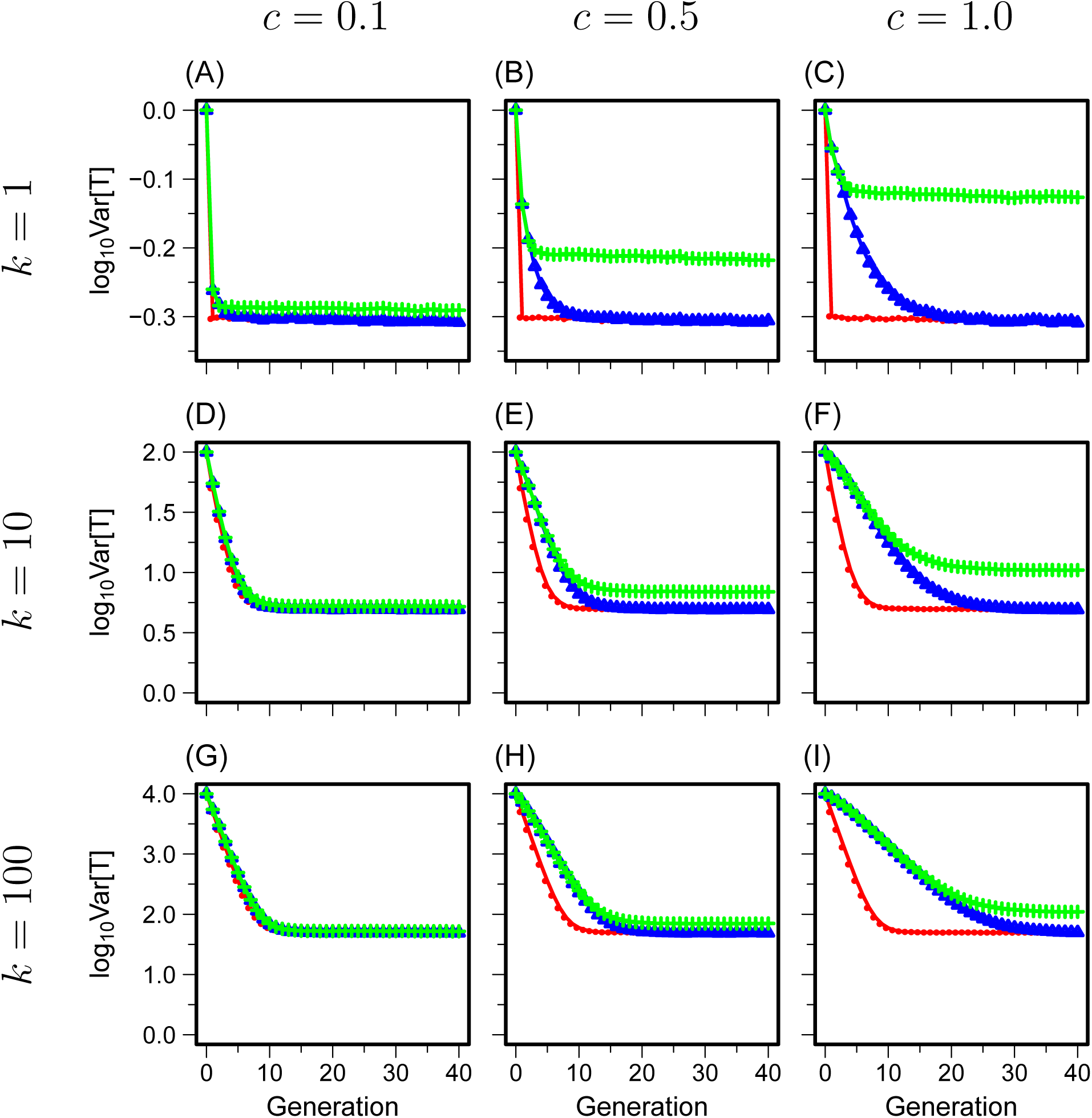
Variance of the phenotype (Var[*T*]) as a function of time. The simulations shown are the same ones from Fig. 4. (A) *k* = 1, *c* = 0.1. (B) *k* = 1, *c* = 0.5. (C) *k* = 1, *c* = 1.0. (D) *k* = 10, *c* = 0.1. (E) *k* = 10, *c* = 0.5. (F) *k* = 10, *c* = 1.0. (G) *k* = 100, *c* = 0.1. (H) *k* = 100, *c* = 0.5. (I) *k* = 100, *c* = 1.0. Colors and symbols follow Figure 4. The y-axis is plotted on a logarithmic scale.

Having examined the behavior of Cor[*H*_*A*_, *T*], Var[*H*_*A*_], and Var[*T*] in the base case, we now explore the effect of the assortative mating strength *c* and the parameters involving the quantitative trait—the number of loci *k*, allele frequencies *p*_*i*_ and *q*_*i*_, and trait contribution at each locus *X*_*i*_—on these quantities.

### 4.2 Assortative Mating Strength (*c*)

#### 4.2.1 Cor[*H*_*A*_, *T*]

Each row of Figure 4 illustrates the influence of the assortative mating strength *c* on Cor[*H*_*A*_, *T*] with a fixed number of trait loci *k*, and each column depicts the effect of the number of loci *k* on Cor[*H*_*A*_, *T*] with fixed assortative mating strength *c*. All parameters other than *c* and *k* are held constant at the base case values.

With different assortative mating strengths and numbers of trait loci, *c* = 0.1, 0.5, 1.0 and *k* = 1, 10, 100, the qualitative behavior of Cor[*H*_*A*_, *T*] over time remains the same as in the base case. As before, we observe decay in Cor[*H*_*A*_, *T*] under all three mating models, with random mating decoupling ancestry and trait values the most rapidly. Cor[*H*_*A*_, *T*] remains higher for longer under assortative mating by phenotype than under assortative mating by admixture fraction. The rate of decay and the degree to which the patterns differ across the three mating models depend on the assortative mating strength and the number of loci.

If assortative mating is weak (*c* = 0.1 in Figure 4A, D, G), then the effect of assortative mating is small, and thus, Cor[*H*_*A*_, *T*] under assortative mating by admixture and by phenotype closely follows that under random mating. This pattern is seen irrespective of the number of loci. Note that in the limit of *c* = 0, the assortative mating and random mating models are identical because the mating function in Eq. 4 becomes a constant, the same value for all three mating models.

Comparing panels within rows of Figure 4, the results from random mating are identical, as the assortative mating strength does not affect the random mating model. Under both assortative mating models, however, Cor[*H*_*A*_, *T*] increases with the assortative mating strength. In an extreme case of complete assortment (*c* → ∞), the correlation would stay constant at 1 across all generations: we start with complete correlation between admixture and phenotype, and *c* → ∞ implies that only identical individuals can mate, so that the correlation persists unchanged.

The difference among the three models increases with the assortative mating strength given a fixed number of trait loci. The difference is the greatest if *k* = 1 and *c* = 1.0 (Figure 4C). Even after 40 generations, assortative mating by trait retains a high correlation at 0.788, whereas the corresponding values under random mating and assortative mating by admixture are 0.006 and 0.010, respectively.

#### 4.2.2 Var[*H*_*A*_] and Var[*T*]

The plots of Var[*H*_*A*_] in Figure 5 and Var[*T*] in Figure 6 consider the same simulations that appear for Cor[*H*_*A*_, *T*] in Figure 4. As is seen in classical work [26, 35, 36], compared to random mating, assortative mating increases the variance of the property on which assortment takes place. Thus, the variance of the admixture fraction is increased to a greater extent under mating by admixture fraction than under mating by phenotype. Similarly, the variance of the phenotype is increased to a greater extent under mating by phenotype than under mating by admixture fraction. Both types of assortative mating increase both Var[*H*_*A*_] and Var[*T*] compared with random mating.

The variance-increasing effect of the assortative mating is visible when comparing panels within each row. For low assortative mating strength (*c* = 0.1), panels A, D, and G in Figures 5 and 6 show that minimal differences in Var[*H*_*A*_] and Var[*T*] exist between mating models. As *c* increases, for a given number of loci, Figures 5 and 6 display increased differences between random and assortative mating, with maximal separation at the largest assortative mating strength simulated, *c* = 1 (panels C, F, I). The random mating model is unaffected by the assortative mating strength *c*, as was seen with Cor[*H*_*A*_, *T*] in Section 4.2.1.

### 4.3 Number of Trait Loci (*k*)

#### 4.3.1 Cor[*H*_*A*_, *T*]

A comparison of panels within columns of Figure 4 shows that under random mating, with more loci associated with the phenotype, the ancestry-phenotype correlation is higher and stays high for longer. In other words, it takes longer for *H*_*A*_ and *T* to become decoupled. In particular, under random mating, the correlation between trait and ancestry falls below 0.5 at *g* = 3 if *k* = 1, *g* = 6 if *k* = 10, and *g* = 10 if *k* = 100, independent of the assortative mating strength.

As the number of loci increases, results from the models with assortative mating by phenotype and by admixture become similar. If (*c, k*) = (1, 100) (Figure 4I), then it takes 24 generations for Cor[*H*_*A*_, *T*] values under the two models to differ by more than 0.1. Corresponding times for (*c, k*) = (1, 1) (Figure 4C) and (*c, k*) = (1, 10) (Figure 4F) are *g* = 6 and *g* = 15, respectively. Recall that the admixture fraction represents the probability that a random allele at a random autosomal genetic locus originates from source population *S*_1_, assuming infinitely many loci. In the *k* → ∞ limit, with the whole genome contributing to the trait, the assortative mating models by admixture and by phenotype would behave in exactly the same way.

#### 4.3.2 Var[*H*_*A*_] and Var[*T*]

Comparing panels within columns in Figure 5, for a given assortative mating strength, Var[*H*_*A*_] under assortative mating by admixture follows the same curve irrespective of the number of loci. Because the mating probability is independent of trait values if mating assortatively by admixture, *k* has no effect.

As in the base case (Section 4.1.2), both assortative mating models have higher Var[*H*_*A*_] and Var[*T*] than random mating. Of the two assortative mating models, assortative mating by admixture fraction has greater Var[*H*_*A*_] than assortative mating by trait at each generation. For Var[*T*], assortative mating by trait has greater values than assortative mating by admixture fraction. As was seen with Cor[*H*_*A*_, *T*] (Section 4.3.1), for Var[*H*_*A*_] and Var[*T*], the difference between random mating and both assortative mating models increases with *k*, and the difference between the two assortative mating models diminishes as *k* increases.

### 4.4 Allele Frequencies (*p*_*i*_ and *q*_*i*_)

Departing further from the base case, we next evaluate the effect of the allele frequencies, *p*_*i*_ and *q*_*i*_, on the quantities of interest. Instead of treating the two source populations as fixed for different alleles, the frequencies *p*_*i*_ and *q*_*i*_ are now sampled according to the simulation procedure described in Section 3.1. Because our results show a monotonic trend across the number of loci we examined (Section 4.3), we focus this analysis on a single value of *k* = 10, the number of loci corresponding to the base case.

#### 4.4.1 Cor[*H*_*A*_, *T*]

Figure 7A displays Cor[*H*_*A*_, *T*] under the model with simulated rather than fixed allele frequencies. Cor[*H*_*A*_, *T*] starts from a lower correlation value at time *g* = 0, 0.456, compared to the base case (Figure 4E) value of 1. If all loci have *X*_*i*_ = 1, as shown in Eq. 1, then an individual’s trait value is determined by the number of “1” alleles across the trait loci. Because the allele “1” is randomly drawn at each locus *i* = 1, 2, *…, k* with probabilities *P* (*L*_*ij*_ = 1 | *M* = *S*_1_) = *p*_*i*_ and *P* (*L*_*ij*_ = 1 | *M* = *S*_2_) = *q*_*i*_ with *j* = 1, 2 (Section 2.2) and the mean absolute difference between simulated *p*_*i*_ and *q*_*i*_ across *k* loci is small, some individuals in the source population *S*_1_ have lower trait values than some individuals in *S*_2_, and vice versa. However, due to the constraint *p*_*i*_ ≥ *q*_*i*_ across all trait loci, individuals from *S*_1_ have higher probability of having a larger trait value than those from *S*_2_. This property accounts for the nonzero correlation between ancestry and trait present in the source populations outside the base case setting.

**Figure 7:**
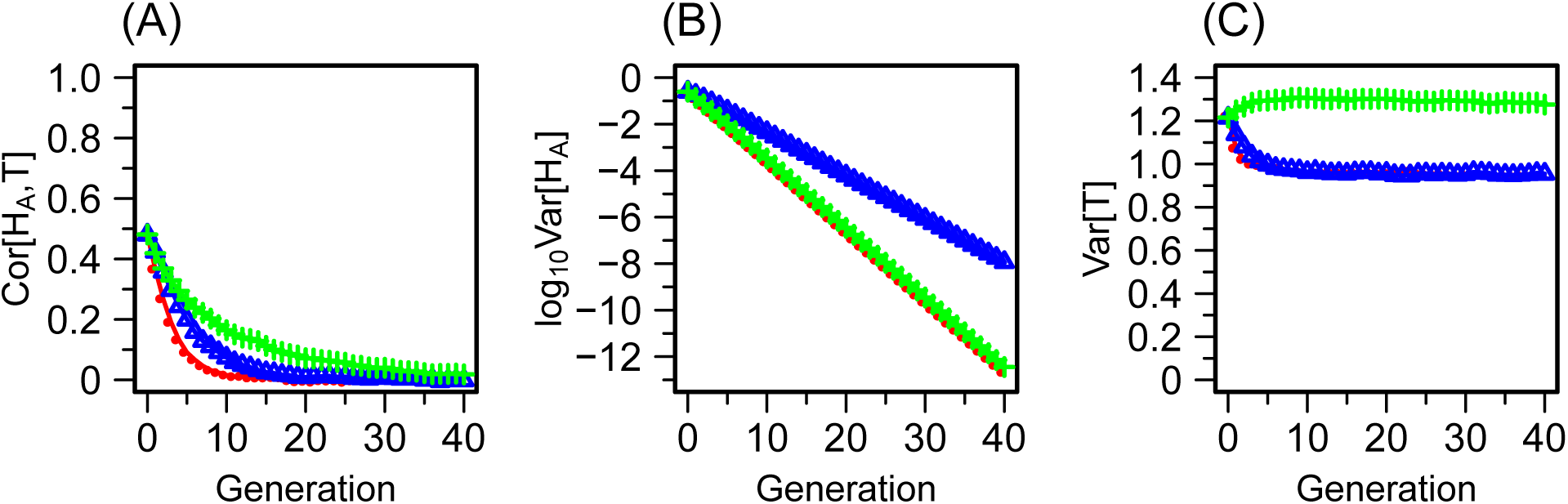
Cor[*H*_*A*_, *T*], Var[*H*_*A*_], and Var[*T*] in a model in which the allele frequencies *p*_*i*_ and *q*_*i*_ in the source populations *S*_1_ and *S*_2_ are drawn from a simulation rather than being treated as fixed at 1 and 0, respectively. All other parameters are kept at the values of the base case (Section 3.2). (A) Correlation between admixture fraction and trait (Cor[*H*_*A*_, *T*]). (B) Variance of the admixture fraction (Var[*H*_*A*_]). (C) Variance of the phenotype (Var[*T*]). Colors and symbols follow Figure 4. The figure relies on a single replicate of simulated allele frequencies *p*_*i*_ and *q*_*i*_ following a genetic drift model in which *S*_1_ and *S*_2_ descend from a common ancestral population, as described in Section 3.1. The simulated allele frequencies across *k* = 10 loci have mean values of 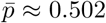 and 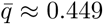 and variance 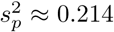 and 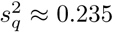. If we let *δ*_*i*_ = *p*_*i*_ −*q*_*i*_, with *δ*_*i*_ > 0, then the mean of the allele frequency difference across the 10 loci is 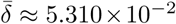, with 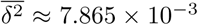. Across *k* = 10 loci, *F*_*ST*_ ≈ 0.075, as computed using Eq. 14 of [22]. The y-axis of Var[*H*_*A*_] is plotted on a logarithmic scale. Results using other replicates of simulated allele frequencies with *k* = 10 are shown in Figure S4.

The qualitative differences between the three mating models remain similar to the base case, as shown in Figure 7A. All three mating models, however, show an increased rate of decoupling between the admixture fraction and the trait, in that the correlation decreases more rapidly. For random mating, it takes only 4 generations for Cor[*H*_*A*_, *T*] to drop to below half of its starting value, reaching 0.170. The corresponding values under assortative mating by admixture fraction and assortative mating by trait are *g* = 5 (Cor[*H*_*A*_, *T*] = 0.240) and *g* = 10 (Cor[*H*_*A*_, *T*] = 0.220), respectively. Compared to the base case, the correlation between ancestry and trait in the source population is weaker if the allele frequencies are drawn from the simulation, and thus, the “1” allele does not necessarily trace back to the source population *S*_1_. Under this setting, the effect of the Mendelian noise in decoupling of admixture fraction and phenotype in producing admixed individuals becomes more significant than the base case.

#### 4.4.2 Var[*H*_*A*_] and Var[*T*]

The admixture fraction values at the source populations are not affected by the allele frequencies: *H*_*A*_ = 1 and *H*_*A*_ = 0 for all individuals in *S*_1_ and *S*_2_, respectively. If *s*_1,0_ = *s*_2,0_ = 0.5, then Var[*H*_*A*_] starts at 0.25 in the founding parental pool, irrespective of the allele frequencies. Comparing Figure 7B and 5E, the Var[*H*_*A*_] curves under random mating (red) and assortative mating by admixture (blue) are not affected by the change in allele frequencies *p*_*i*_ and *q*_*i*_, holding other parameters fixed. Under random mating and assortative mating by admixture, mate choice is independent of the parameters that affect the quantitative trait, and thus, the change in *p*_*i*_ and *q*_*i*_ does not alter the admixture fraction distribution at each generation.

By contrast, under assortative mating by phenotype, Var[*H*_*A*_] (green) is affected by the change in the nature of the allele frequencies. Var[*H*_*A*_] under assortative mating by phenotype closely follows that under random mating. The simulated allele frequencies have relatively small differences 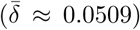 between source populations *S*_1_ and *S*_2_. With *X*_*i*_ = 1 for all loci, the between-group difference in trait values is small as well, whereas all individuals in *S*_1_ and *S*_2_ still have *H*_*A*_ = 1 and *H*_*A*_ = 0, respectively. Therefore, with the simulated allele frequencies, the effect on the admixture fraction of assortative mating by phenotype is similar to that in the random mating case. This scenario contrasts with the base case, where allele “1” can be associated with the source population *S*_1_ with certainty, and Var[*H*_*A*_] under assortative mating by phenotype behaves similarly to the case of assortative mating by admixture fraction.

With the simulated allele frequencies, Var[*T*] = 0.877 in the founding parental pool. At *g* = 1, Var[*T*] values under random mating and under assortative mating by admixture are 0.784 and 0.825, respectively. Assortative mating by admixture maintains higher Var[*T*] than random mating until *g* = 8 and then follows the Var[*T*] curve for random mating. By contrast, Var[*T*] under assortative mating by trait gradually increases until *g* = 13, at which it achieves its maximum of 0.977, and then decreases to 0.935 at *g* = 40.

### 4.5 Trait Contributions of Individual Loci (*X*_*i*_)

Returning to the case with fixed allele frequencies of 1 and 0 in the source populations, we next examine the case in which the trait has the property that both alleles have equal probability of being the “+” allele, as described in Section 2.2: 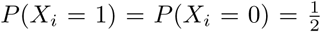 for all *i* = 1, 2, *…, k*. Figure 8 displays the results using the number of trait loci from the base case, *k* = 10. The qualitative behavior of the result does not depend on the number of loci with the other parameters fixed.

**Figure 8:**
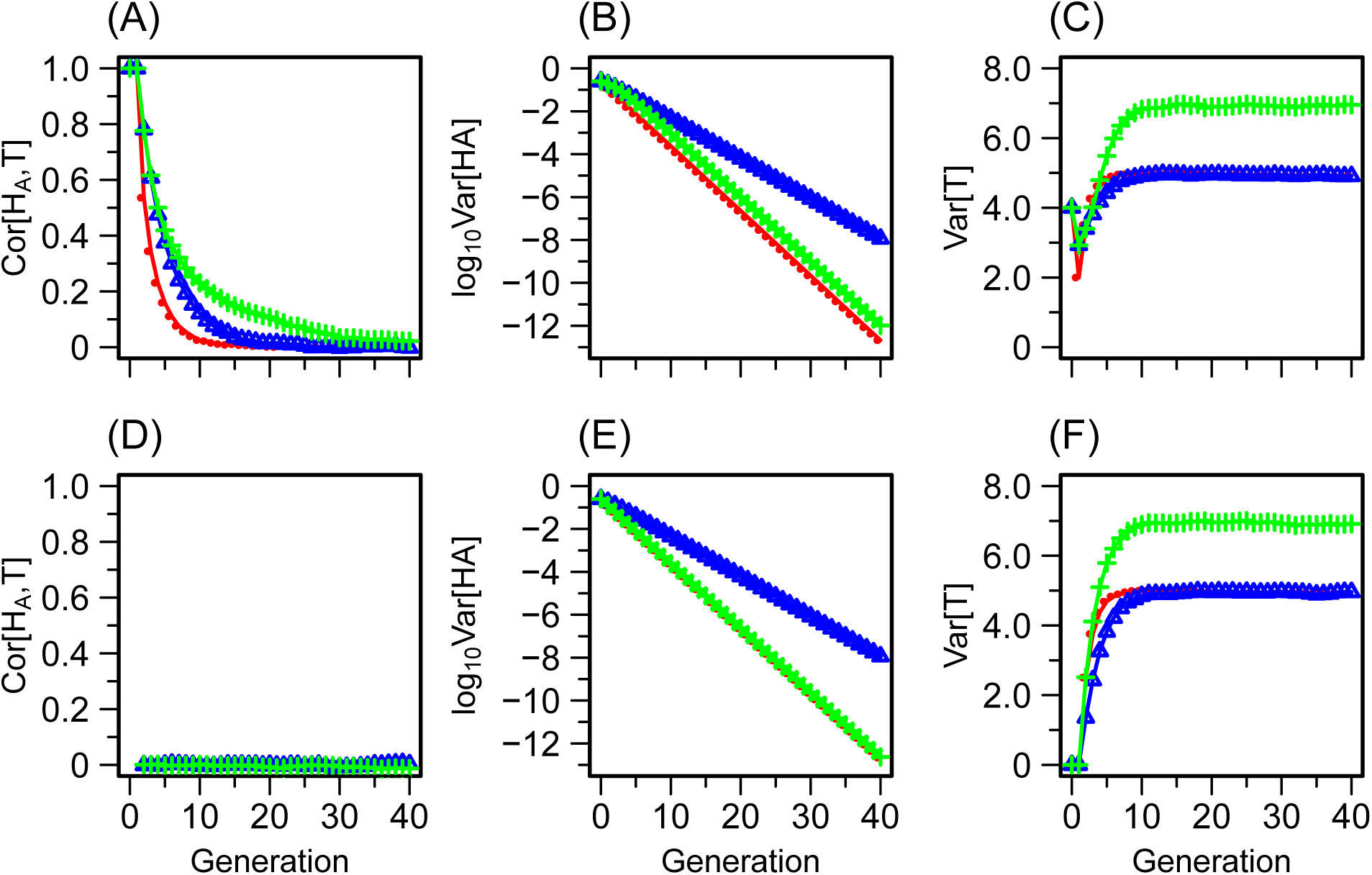
Cor[*H*_*A*_, *T*], Var[*H*_*A*_], and Var[*T*] under a trait model in which trait loci do not systematically have greater values in one source population: *P* (*X*_*i*_ = 1) = *P* (*X*_*i*_ = 0) = 0.5. All other parameters are kept at the values of the base case (Section 3.2). Of *k* = 10 trait loci, we denote the number of randomly selected loci to have *X*_*i*_ = 1 by *z*. (A) Cor[*H*_*A*_, *T*], *z* = 6. (B) Var[*H*_*A*_], *z* = 6. (C) Var[*T*], *z* = 6. (D) Cor[*H*_*A*_, *T*], *z* = 5. (E) Var[*H*_*A*_], *z* = 5. (F) Var[*T*], *z* = 5. Colors and symbols follow Figure 4. Panels A-C and D-F each relies on a single replicate of a set of *X*_*i*_ obtained by sampling the *X*_*i*_ from a Binomial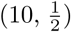 distribution and retaining those with the specified value of *z* = 6 (top panels) and *z* = 5 (bottom panels). The y-axis of Var[*H*_*A*_] is plotted on a logarithmic scale. Results using other replicates of simulated allele frequencies with *k* = 10 are shown in Figures S5 (for *z* = 6) and S6 (for *z* = 5).

#### 4.5.1 Cor[*H*_*A*_, *T*]

If we let the number of loci with *X*_*i*_ = 1 be *z*, then the number of loci with *X*_*i*_ = 0 is *k* − *z*. Because *p*_*i*_ = 1 and *q*_*i*_ = 0 across all loci in the base case, the trait value is 2*z* for every individual in *S*_1_ and 2(*k* − *z*) for every individual in *S*_2_ (Eq. 1). For a randomly generated set of *X*_*i*_, {*i* = 1, 2, *…, k*}, under 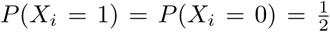, if *z* ≠ *k* − *z*, then Cor[*H*_*A*_, *T*] = 1 in the founding parental pool 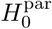, as shown in Figure 8A. However, compared with the base case (Figure 3.2), the correlation decays much more rapidly. With the *P* (*X*_*i*_ = 1) ≠ 1 setting, the ancestry and trait are not as tightly coupled in the source populations. However, as in Section 4.1-4.4, assortative mating by phenotype preserves the correlation for the longest, and random mating decouples the correlation the fastest of the three mating models.

If the numbers of loci with *X*_*i*_ = 1 and *X*_*i*_ = 0 are equal (*z* = *k* − *z*), then all individuals in the source populations have trait value *k* irrespective of their origin, and thus, no correlation exists between trait and ancestry in the source population. Hence, Cor[*H*_*A*_, *T*] is 0 in the founding parental pool 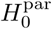, and the correlation remains at 0 throughout the time simulated, irrespective of the mating type (Figure 8D).

#### 4.5.2 Var[*H*_*A*_] and Var[*T*]

The panels of Figure 8 display results under 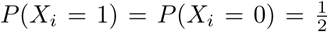, fixing other parameters as in base case. By the same reasoning as in Section 4.4.2, the change in parameters involving the quantitative trait does not affect Var[*H*_*A*_] under random mating and under assortative mating by admixture fraction. A comparison of Figure 8B and 8E with Figure 5E shows that Var[*H*_*A*_] values under assortative mating by admixture fraction are not affected by the change in the *X*_*i*_.

For *z* ≠ *k* − *z*, results with *z* = 6 and *k* − *z* = 4 appear in Figure 8B. Some correlation between the admixture fraction and allele “1” exists in the source populations, and thus, Var[*H*_*A*_] for assortative mating by trait (green) somewhat follows that for assortative mating by admixture (blue). However, compared to the base case (Figure 5E), where the “1” allele can be traced back to *S*_1_ with certainty, a more noticeable deviation from the blue curve is observed. As *z* increases from 6 to 10, the pattern is similar; the quantitative behavior of Var[*H*_*A*_] would approach the base case, equivalent to *z* = 10 (green curve in Figure 5E).

In the founding parental pool, Var[*T*] = 4.002 in our example with *z* ≠ *k* − *z*. After one generation of mating, Var[*T*] drops to 1.997, 2.934, and 2.923, under random mating, assortative mating by admixture fraction, and assortative mating by trait, respectively. From *g* = 2, Var[*T*] gradually increases and achieves steady state values for the three models near 4.942 at *g* = 5, 4.934 *g* = 8, and 6.929 at *g* = 14, respectively. In accord with Section 4.1-4.4, assortative mating by trait has the highest Var[*T*] values across generations.

If *z* = *k* − *z* (Figure 8E), then allele “1” has equal probability of traced back to either source population. In this scenario, the Var[*H*_*A*_] curve from assortative mating follows the Var[*H*_*A*_] curve from random mating. Because all individuals have the same trait value in the founding parental pool irrespective of their origin, Var[*T*] = 0 at *g* = 0. For all three mating models, Var[*T*] gradually increases from *g* = 1 to achieve steady state values that are the same as those from the *z* ≠ *k* − *z* case.

## 5 Discussion

In this paper, we have devised a mechanistic admixture model in which an admixed population is formed from contributions of a pair of mutually isolated source populations under assortative mating, either by admixture level or by a quantitative phenotype. The approach includes a quantitative-genetic model that relates a quantitative phenotype to underlying loci affecting its trait value and, ultimately, to mate choices. The admixture level and the quantitative phenotype are studied using a discrete-time recursion that describes the evolution of the admixed population. Under this model, we have examined the correlation between genetic ancestry and phenotype in the admixed population as a function of time under three mating models: random mating, assortative mating by admixture fraction, and assortative mating by phenotype.

Initially, ancestry and phenotype are coupled, as the source populations differ in phenotype. Random mating then decouples the correlation between ancestry and trait faster than is seen in both assortative mating models (Figure 4), and assortative mating by phenotype maintains the correlation to a greater extent than does assortative mating by admixture (Figure 4). Compared with random mating, in a similar manner to classic assortative mating models [26, 35, 36], the assortative mating increases the population variance of the property on which the assortment is based (Figures 5 and 6). In fact, our Eq. 13 multiplies the variance of admixture in a model without assortment [1] by a factor that increases with positive assortative mating.

Increasing the strength of assortative mating magnifies the difference observed among the models in the speed at which the correlation declines (Figure 4). Generally similar qualitative patterns are observed if the source populations are regarded as having fixed differences (Figure 4) or merely frequency differences in alleles affecting the quantitative trait (Figures 7 and 8).

A key observation is that, as the number of loci underlying the quantitative trait increases, the difference in trajectories between the two assortative mating models decreases. Because assortative mating by admixture fraction affects all loci, whereas assortative mating by trait affects only trait loci and their genomic neighbors, as the trait is determined by an increasing number of loci, the behavior of assortative mating by trait increasingly follows that of assortative mating by ancestry (Figure 4). As the number of loci affecting the phenotype increases, assortative mating by trait increasingly reflects assortative mating by admixture, and the correlation between ancestry and trait value persists for longer (Figure 4).

Our study was motivated partly by a hypothesis of Parra et al. [8] claiming that assortative mating by the *color* phenotype in Brazil could eventually decouple *color* from genetic ancestry, so that largely separate subpopulations with distinct *color* could eventually possess similar levels of African genetic ancestry. We have seen not only that assortative mating by a quantitative trait that differs between source populations can decouple the phenotype from the genetic ancestry, but that random mating can decouple the phenotype from genetic ancestry as well. Moreover, the decoupling is *slower* with assortative mating than it is with random mating. In an admixed population with assortative mating that is heavily influenced by a salient phenotype (such as *color* in the scenario of Parra et al. [8]), mating by other genetically influenced phenotypes is random, or perhaps less strongly assortative. Thus, in an admixed population, we might expect that among all the traits to which genotypes contribute, traits that have little influence on mating behavior will decouple from ancestry most rapidly. Those traits on which assortment does occur, such as *color* in Brazil, will be the slowest to decouple from ancestry—but under our model, they eventually will do so. Thus, in an admixed population, phenotypes that once reflected ancestry in the source populations might no longer be predictive of genetic ancestry after a sufficient length of time has passed.

The focus of our simulations has been on understanding demographic phenomena, but the model is relevant to efforts to investigate determinants of disease traits in admixed populations. For example, in admixture-mapping studies and in studies of health disparities involving admixed populations, correlations of phenotypes and admixture levels are often computed [37–40]. The mechanistic model can potentially provide insights into the way in which these correlations change over time in scenarios in which specific trait architectures are of interest.

We note that we have examined a model with a single admixture event at the founding of the admixed population. Under this idealized model, even if the founding admixed population starts with a perfect correlation between admixture fraction and trait value, the correlation decreases over time and eventually approaches zero in the absence of further influx from source populations. In principle, the framework can account for continuous influx from the source populations. If ancestry–trait correlation exists in source populations, then such influx would be expected to slow the decoupling between admixture and phenotype in the admixed populations under all three mating models, while qualitatively maintaining their relative order in the rate of decoupling.

Although the theoretical framework we have developed can incorporate various genetic architectures and population admixture processes (including disassortative mating), our model has a number of limitations. First, it does not include sex bias during the admixture event, a phenomenon that often occurs in admixture processes [17, 41, 42]. A recent genetic model in a scenario with continuing contributions from the sources does allow for sex bias with assortative mating, but with no phenotype and with the assortative mating occurring by population membership—in the admixed population or in one of the sources—rather than by the admixture level itself [28]. Next, we have chosen to model admixture in an individual as the mean of parental admixture levels; this approach does not account for stochasticity during genetic transmission. Our quantitative trait model does not incorporate dominance, spatial positioning of trait loci along a genome, epistasis, environmental effects on the phenotype, or genotype-by-environment interaction. The latter pair of limitations might be particularly important in using the results in human data, as numerous studies have shown that assortative mating often operates on sociocultural traits [43–46]. It will be important to extend our theoretical framework to include additional features of the quantitative-genetic model, such as by incorporating dominance, linkage, variable effect sizes across the loci contributing to the quantitative trait, and varying heritability of the phenotype.\

## Acknowledgments

We acknowledge support from National Institutes of Health grants R01 HG005855 and F32 GM130050 and National Science Foundation grant BCS-1515127.

## Supplementary Material

**Figure S1:**
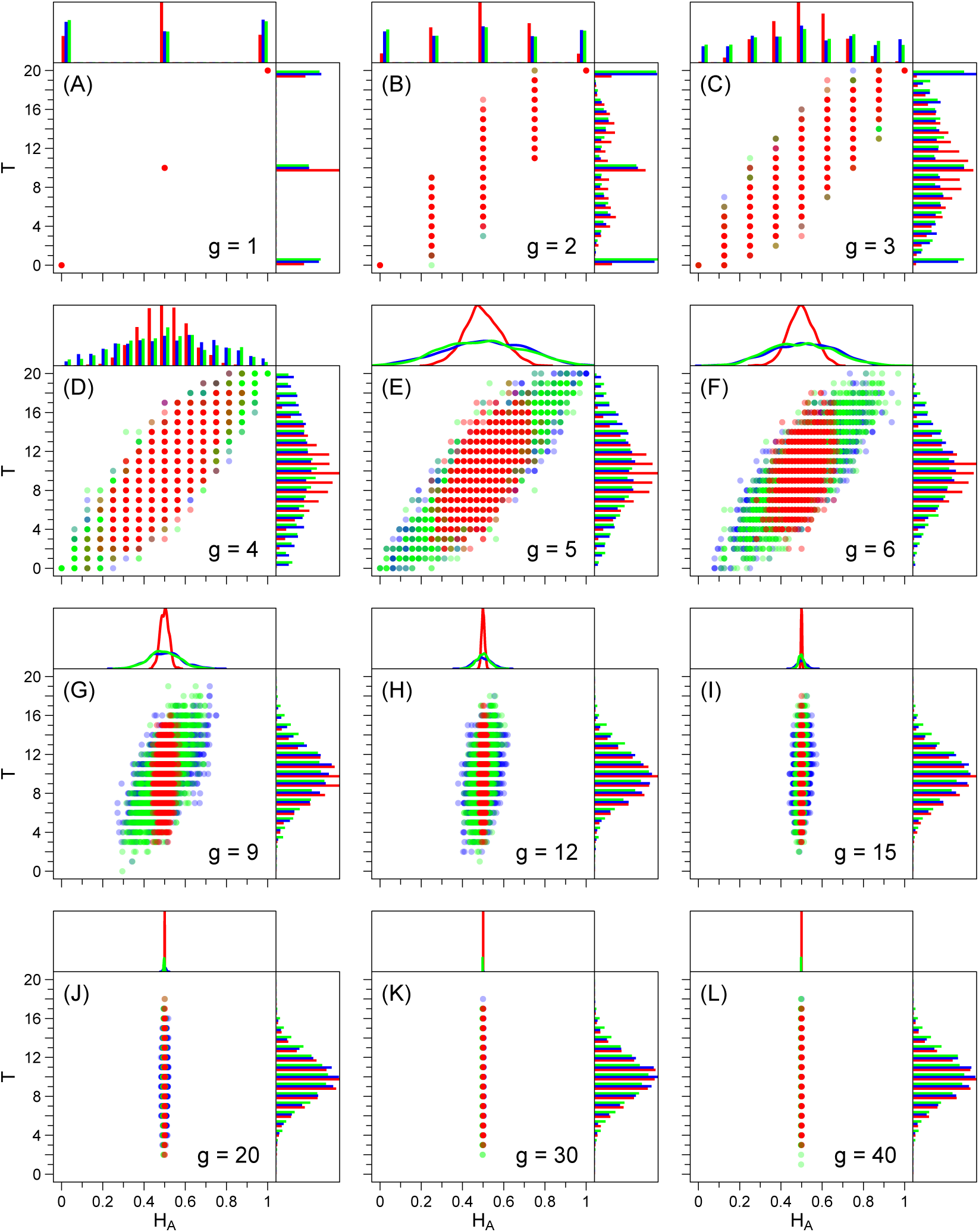
Joint distribution of *H*_*A*_ and *T* as a function of time. The simulation shown is the same one from Figures 4E, 5E, and 6E, using the parameters from base case (Section 3.2). As described in Section 2.1 and 2.2, the possible values for the admixture fraction at generation *g* are 0, 1*/*2^*g*^, 2*/*2^*g*^, *…*, (2^*g*^ − 1)*/*2^*g*^, 1, whereas the possible values for the trait are 0, 1, *…*, 2*k* across all generations. In each panel, the top, right, and center plots display a marginal distribution of *H*_*A*_, a marginal distribution of *T*, and a joint distribution of *H*_*A*_ and *T*, respectively. Colors follow Figure 4.

**Figure S2:**
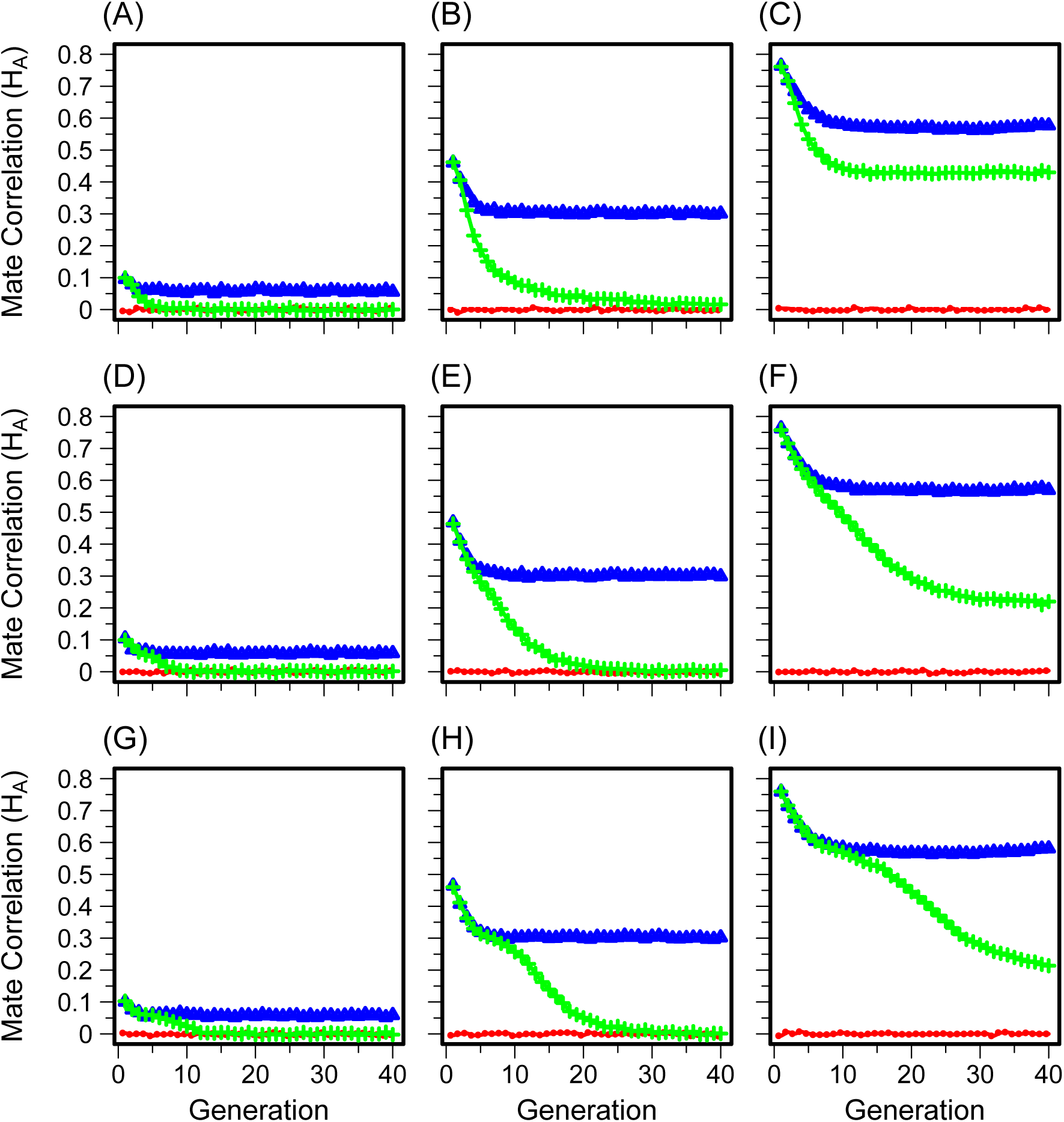
The correlation coefficient 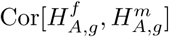 between the admixture fractions of the members of mating pairs as a function of time. The simulations shown are the same ones from Fig. 4. (A) *k* = 1, *c* = 0.1. (B) *k* = 1, *c* = 0.5. (C) *k* = 1, *c* = 1.0. (D) *k* = 10, *c* = 0.1. (E) *k* = 10, *c* = 0.5. (F) *k* = 10, *c* = 1.0. (G) *k* = 100, *c* = 0.1. (H) *k* = 100, *c* = 0.5. (I) *k* = 100, *c* = 1.0. Colors and symbols follow Figure 4.

**Figure S3:**
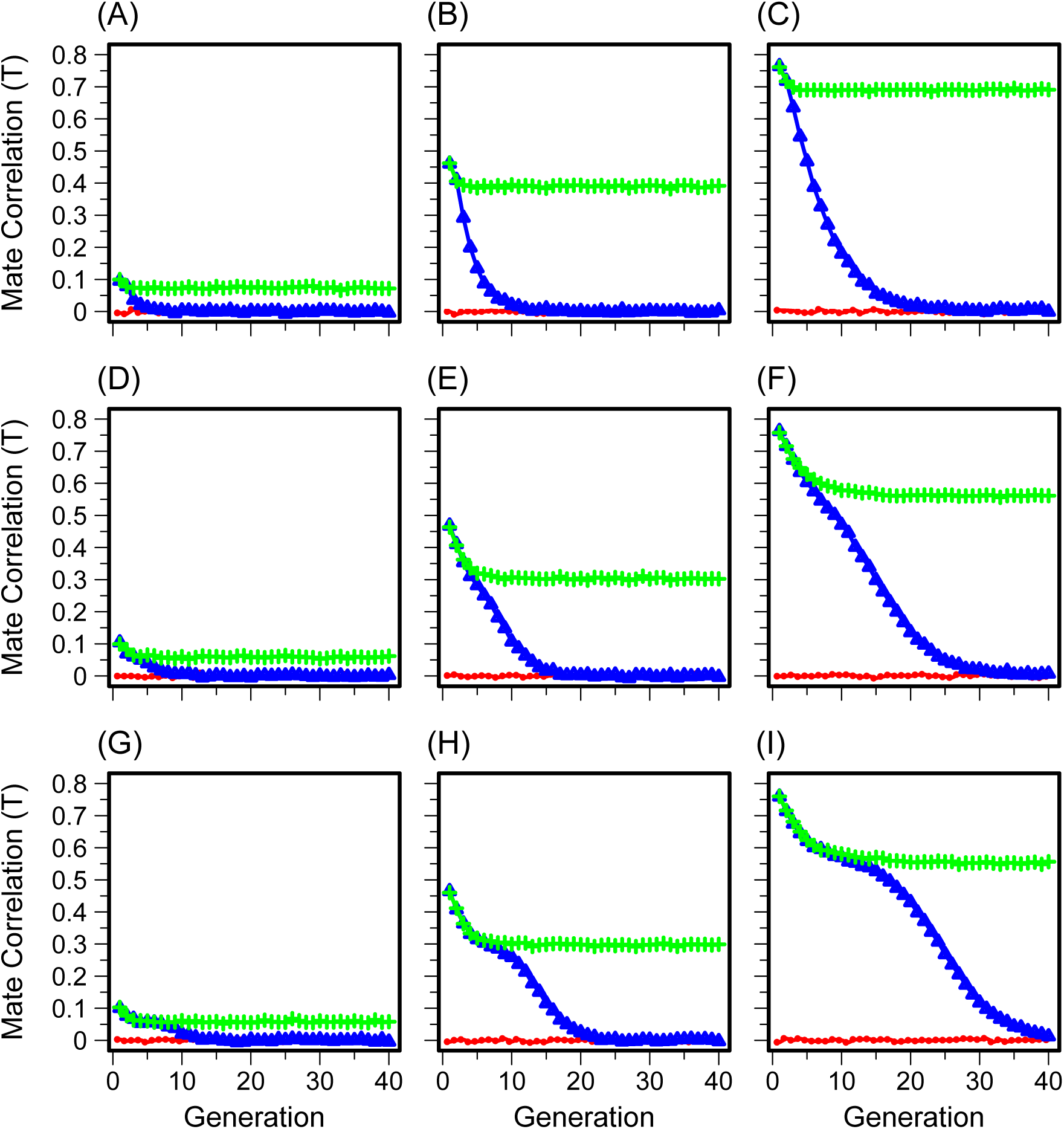
The correlation coefficient 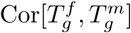 between the phenotypes of the members of mating pairs as a function of time. The simulations shown are the same ones from Fig. 4. (A) *k* = 1, *c* = 0.1. (B) *k* = 1, *c* = 0.5. (C) *k* = 1, *c* = 1.0. (D) *k* = 10, *c* = 0.1. (E) *k* = 10, *c* = 0.5. (F) *k* = 10, *c* = 1.0. (G) *k* = 100, *c* = 0.1. (H) *k* = 100, *c* = 0.5. (I) *k* = 100, *c* = 1.0. Colors and symbols follow Figure 4.

**Figure S4:**
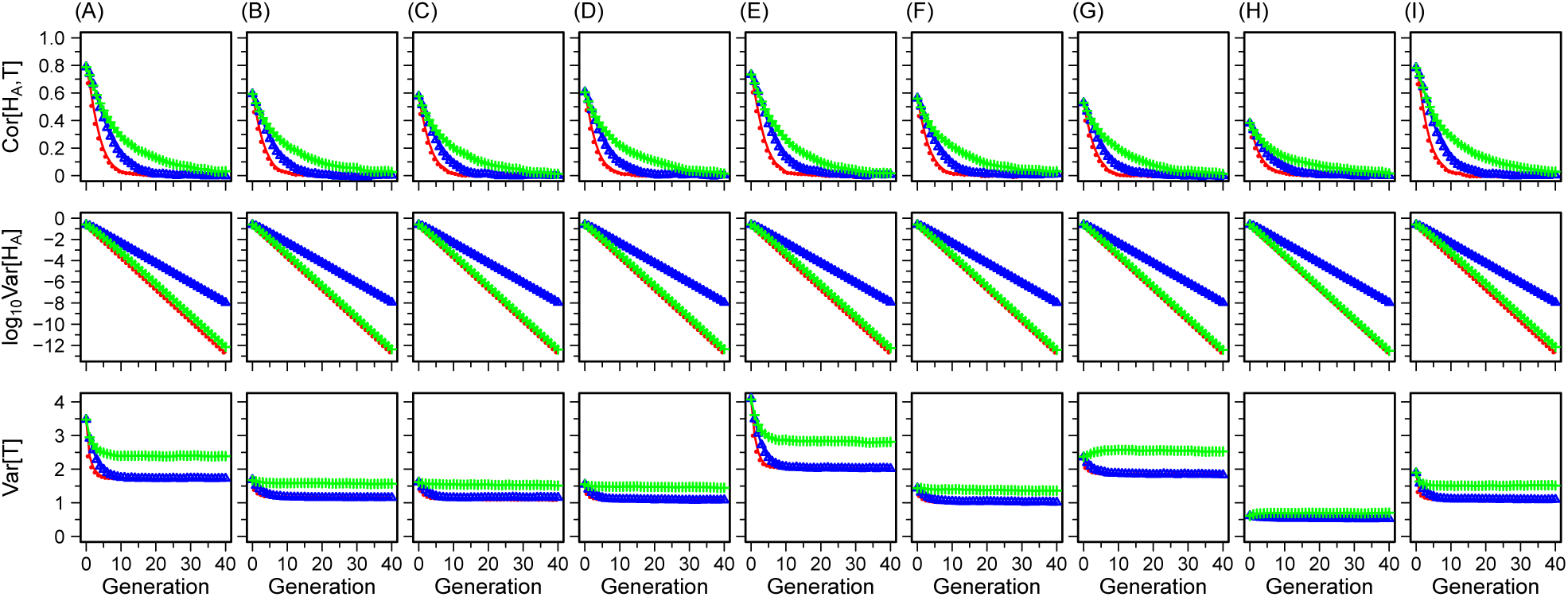
Cor[*H*_*A*_, *T*] (top row), Var[*H*_*A*_] (middle row), and Var[*T*] (bottom row) using different replicate sets of simulated allele frequencies *p*_*i*_ and *q*_*i*_ with *k* = 10 loci, as described in Section 3.1 and Figure 7. Different columns represent results from different replicates of simulated *p*_*i*_ and *q*_*i*_. Colors and symbols follow Figure 4. The y-axis of Var[*H*_*A*_] is plotted on a logarithmic scale with base 10.

**Figure S5:**
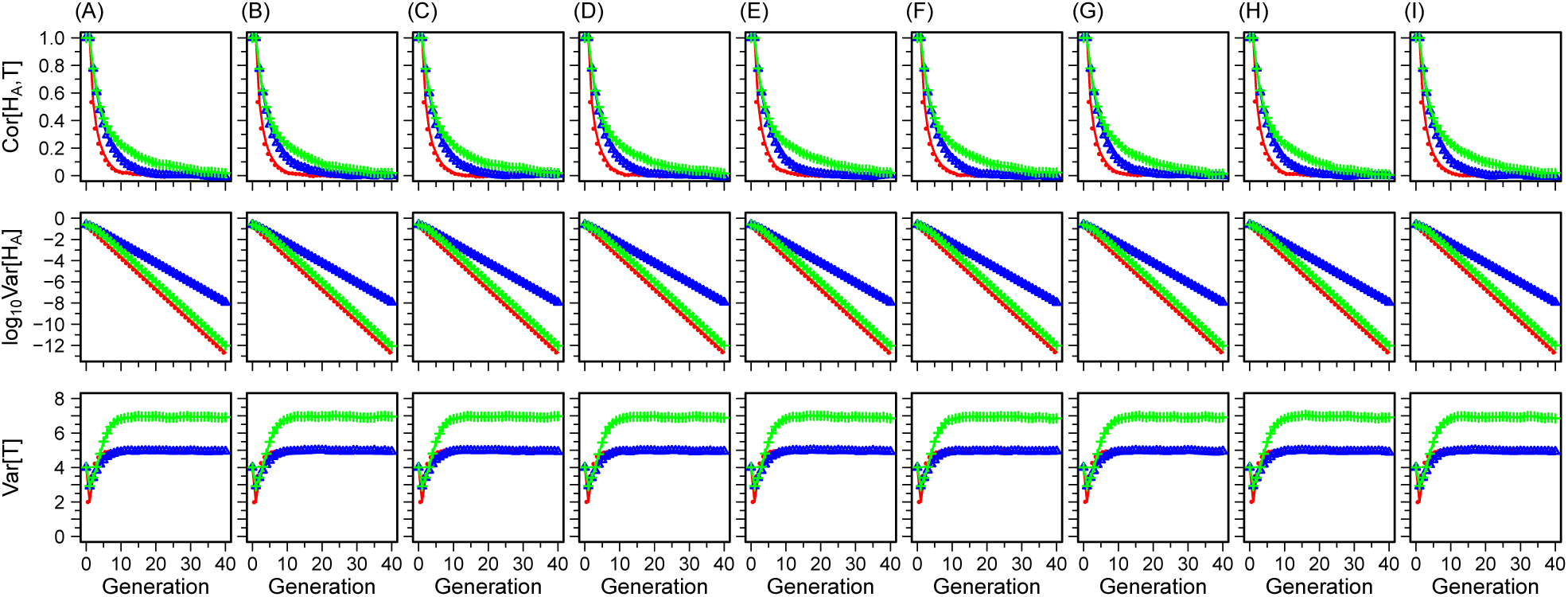
Cor[*H*_*A*_, *T*] (top row), Var[*H*_*A*_] (middle row), and Var[*T*] (bottom row) using different replicates of a set of *X*_*i*_ for *k* = 10 loci sampled from a Binomial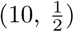 distribution with a constraint *z* = 6 (Section 2.2 and Figure 8A-C). Colors and symbols follow Figure 4. The y-axis of Var[*H*_*A*_] is plotted on a logarithmic scale.

**Figure S6:**
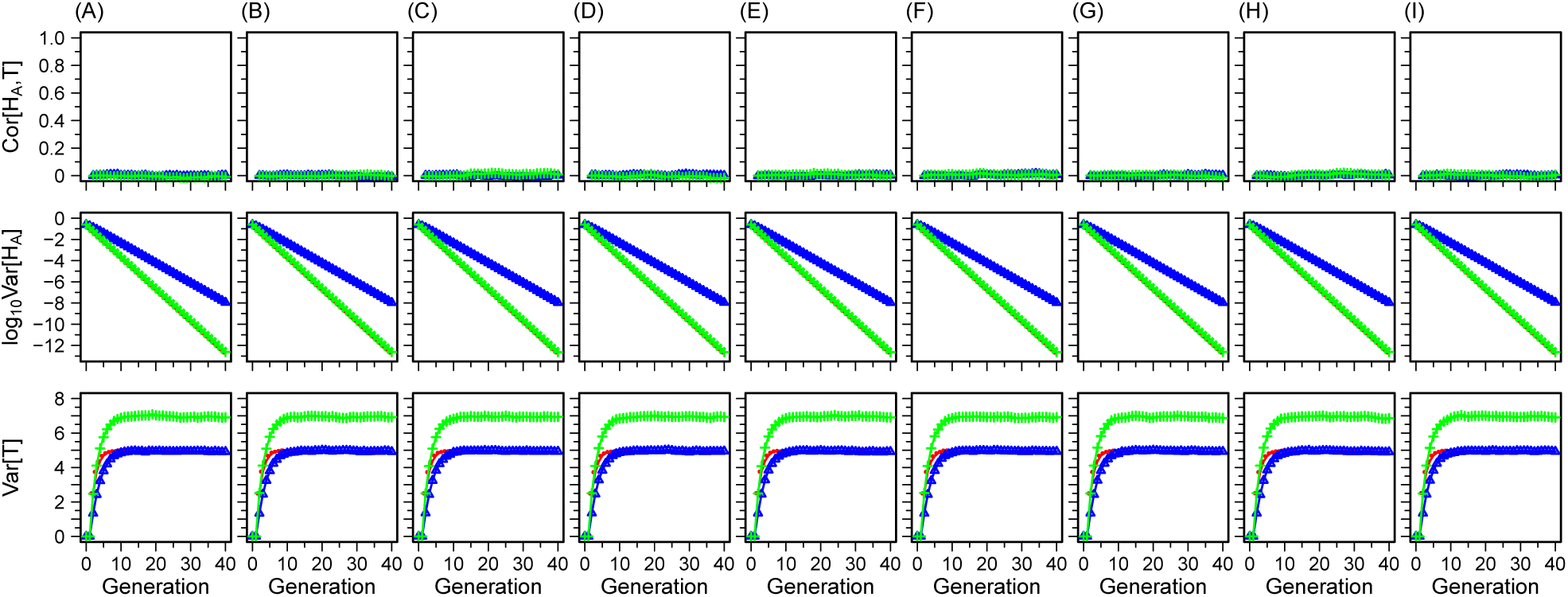
Cor[*H*_*A*_, *T*] (top row), Var[*H*_*A*_] (middle row), and Var[*T*] (bottom row) using different replicates of a set of *X*_*i*_ for *k* = 10 loci sampled from a Binomial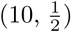 distribution with a constraint *z* = 5 (Section 2.2 and Figure 8D-F). Colors and symbols follow Figure 4. The y-axis of Var[*H*_*A*_] is plotted on a logarithmic scale.

